# Health or disease – a question of rhizomicrobial ecology? The case of Grapevine Trunk Disease

**DOI:** 10.1101/2025.09.02.673213

**Authors:** Islam M. Khattab, Tyra Magold, Florian Lenk, Gunnar Sturm, Noemi Flubacher, Anne-Kristin Kaster, Peter Nick

## Abstract

**Aims:** The incidence of the apoplectic breakdown associated with grapevine trunk diseases (GTDs) is promoted by climate change, which has become a challenge for viticulture worldwide. Outbreak of these conditional diseases is expected to depend on the rhizomicrobiome. However, the impact of the rhizomicrobiome on grapevine resilience has remained poorly understood, particularly regarding its ecological aspects. This study explores the link between GTDs, the rhizomicrobiome, and soil chemistry in vineyards along the Upper Rhine.

**Methods:** Using amplicon sequencing for both fungal and prokaryotic communities, we show that around half of the fungal rhizosphere community is endowed with pathotrophic potential, independently of the health status of the plant, including seventeen previously reported GTD-associated taxa, predominantly Black Foot Disease.

**Results:** In contrast to fungi, bacterial diversity is shifted depending on the micronutrients Fe, Cu, Mn, and Zn. Moreover, taxa enriched in the rhizosphere of asymptomatic vines, such as *Pseudophialocephala* and *Collarina* for the mycobiome, and *Caulobacter*, *Kitasatospora*, and *Entotheonellaceae* for the bacteriome, showed correlations with soil properties. The most prominent feature associated with disease outbreaks was drastic changes of microbial co-occurrence networks. These were significantly increased in the fungi, especially for GTDs taxa, such as *Fomitoporia, Stereum, Phaeomoniella*, and *Neofusicoccum*. By contrast, there was a depletion of many bacteria and their microbial interactions under disease outbreak such as *Isoptericola, Caulobacter, Rhodomicrobium* and *Thioprofundum*.

**Conclusion:** Thus, likely microbial interactions and not the mere presence of GTDs taxa explains disease outbreak. This finding opens new strategies for sustainable management of GTDs.

## Introduction

Grapevine Trunk Diseases (GTDs) threaten viticulture worldwide, accelerated by the ongoing climate change. In France alone, yield losses in 2016 accumulated to 25%, corresponding to approximately 5000 million US$ (https://www.maladie-du-bois-vigne.fr) deficit. The outbreak of different forms of GTDs such as Botryosphaeria dieback, Esca syndrome, Eutypa dieback, Diaporthe dieback, and black foot disease is associated with a wide range of fungal endophytes of around 174 species (Li et al., 2023a). Unlike classical plant diseases, GTDs do not follow the Koch postulates, meaning that the expression of symptoms is not correlated to pathogen abundance, but rather depend on the condition of the host. For example, *Neofusicoccum parvum*, an aggressive fungus causing Botryosphaeria dieback, was found to switch to the necrotrophic phase when the host faces severe drought stress, provoking accumulation of the monolignol precursor ferulic acid. Increases in steady-state levels of ferulic acid might be interpreted by the fungus as a “plant surrender” signal, driving the fungus to secrete a Fusicoccin A aglycon, which afterwards triggers programmed plant cell death (Khattab et al., 2023). In the absence of ferulic acid, the fungus manipulates the homeostasis between defense and growth of the host by secreting an auxin mimic, 4-hydrophenylacetic acid, interfering with specific branches of phytoalexin synthesis (Flubacher et al., 2023).

Conditional pathogenesis is not limited to GTDs, though. While plant-pathogen interactions are often conceptualized as a battle between two opponents, it is important to consider that this viewpoint represents a reduction of a far more complex reality. In fact, the outcome of this battle depends on numerous environmental factors, including the presence of other microorganisms that can have a major impact on the infection process. For example, resistance of a tomato genotype to soil-borne disease was associated with microbiota differing from those in a susceptible genotype (Kwak et al., 2018). As sessile organisms, plants have evolved to regulate the microbial communities in the rhizosphere (for review see Berendsen et al., 2012). The increasing number of examples, where beneficial soil microbes have been found to help plants to survive under environmental challenges (De Vries et al., 2020; Field et al., 2015; Ren et al., 2019) suggest that the interaction between plants and the so-called rhizomicrobiome might be subject of co-evolution. The model of a mutualistic relationship is also supported by findings, where taxonomic structure and function changes depending on plant developmental stage and stress conditions (Berendsen et al., 2012; Gu et al., 2022), supporting the concept of the rhizomicrobiome acting as a “second genome” for plants because of its pivotal role in promoting plant health and resilience (De Vries et al., 2020; Mendes et al., 2011).

Thus, even for the same host genotype and the same physicochemical soil properties, the result of an encounter of a plant with a pathogen can vary between full breakdown and a mitigation even to a degree that the plant remains asymptomatic, depending on the composition of the rhizomicrobiome (Wei et al., 2019). Conversely, shifts in the rhizosphere microbiome could serve as predictive markers of plant resilience to pathogens (Gu et al., 2022; Wei et al., 2019). Furthermore, enriching the soil with synthetic communities of protective taxa might be used as strategy to suppress disease outbreak as shown for bacterial wilt in tomatoes (Lee et al., 2021).

In the context of GTDs, the interactions of grapevine rhizomicrobiome and GTDs have hardly been investigated. Studying the interplay between the fungi causing GTDs, and the rhizomicrobiome could help to sort out either taxa with biocontrol potential to GTDs, or taxa triggering the GTDs outbreak. For instance, a study in young vineyards in China showed that the relative abundance of GTD fungi was irrelevant to their pathogenesis, while the symptoms of GTDs were more linked to the incidence of *Fusarium spp.* in the rhizosphere (Li et al., 2023b). Likewise, wood microbiome analysis for vineyards of different locations in Greece showed that symptomatic wood harboured more *Acremonium alternatum* and *Kalmusia variispora*, fungi not known as causes of GTD symptoms, while members of the bacterial family *Bacillaceae* were depleted in those symptomatic vines (Fotios et al., 2021). This observation corroborates findings, where a specific member of the *Bacillaceae*, *Bacillus subtilis* PTA-271, was found to exert biocontrol activity *in planta* against *Neofusicoccum parvum*, one of the most aggressive GTD fungi (Trotel-Aziz et al., 2019). This bacterial strain modulated accumulation of transcripts for defence-related genes, including those that are regulated by the major defence hormones, salicylic acid and jasmonates. In addition, a glutathione transferase was activated that was proposed to be involved in the catabolic breakdown of the fungal pathogenicity factors (-)-terremutin and (R)-mellein (Trotel-Aziz et al., 2019). Likewise, the immediate inoculation of *Trichoderma* species to pruning wounds in grapevine inhibited infection progress of *Neofusicoccum parvum* and *Diplodia seriata* and achieved a high degree of plant protection (Pollard-Flamand et al., 2022).

The composition and the function of the soil microbiome is shaped by the physiochemical properties of the soil. Soil acidification significantly reduced the potential of microbial communities to inhibit the infections with the phytopathogenic fungus, *Fusarium* (Li et al., 2023). Here, inoculating healthy plants with microbiomes from acidified soils resulted in a remarkable decrease in their ability to resist infection process as well as a downregulation of sulfur metabolism (Li et al., 2023).. Also, micronutrient availability can modulate richness and diversity of the soil microbiome. A survey of 180 sites in China revealed that the structure and function of soil microbiomes was strongly linked to the metallic micronutrients iron, manganese, copper, and zinc (Dai et al., 2023). Specifically, increased Fe and Zn concentrations correlated with ecosystem productivity, which might be a direct consequence of improved plant-nutrient availability, or an indirect effect from altered microbiome composition and gene activity (Dai et al., 2023a). At least for Fe, direct modulation of microbiome-pathogen interactions was demonstrated for the colonisation of tomatoes by *Ralstonia solanacearum* (Gu et al., 2020). Here, under Fe-limited conditions, the rhizomicrobiome of tomato plants was able to outcompete this bacterial pathogen by secreting siderophores. As a result, its growth was suppressed. When this limitation was removed by supplementing iron, the rhizobiome failed to mitigate the infection with *Ralstonia* (Gu et al., 2020).

In viticulture, even small variations in soil properties or water management can significantly affect yields and flavour of economically relevant varieties, such as Chardonnay, Merlot, and Pinot Noir, a phenomenon traditionally known as *terroir*. Those effects are proposed to be linked with shifts in their commensal microbiome (Gilbert et al., 2014). Taxonomic abundance and diversity of both, soil bacterial and fungal flora, is strongly dependent on cultivation practices (Coller et al., 2019). Additionally, the influence of soil microbiome is not confined to the soil, because bacterial taxa isolated from grapevine foliage were tightly associated with the communities in the soil, suggesting that soil may serve as a reservoir for vine-associated microbial flora (Zarraonaindia et al., 2015).

The interplay between grapevine rhizomicrobiome, soil properties, and GTD outbreak has not been addressed yet. However, filling these knowledge gaps is crucial for developing a sustainable approach for grapevine resilience against GTD. Traditional plant breeding methods in viticulture are time consuming and no longer sufficient to cope with the rapidly progressing climate-borne challenges. Targeting beneficial rhizomicrobiota promoting grapevine resistance might act as a fast and sustainable approach against GTDs. This study employed a microbial-ecological strategy, probing the rhizomicrobiome from symptomatic and asymptomatic vines coming from the same vineyard, and sampling over a transsect of more than ten vineyards differing in soil composition to explore the role of the rhizomicrobiome and soil nutrient dynamics for improved grapevine resilience against trunk diseases. By uncovering key interactions between microbes, nutrients, and GTD symptoms, the study contributes to the development of novel strategies for sustainable management of GTDs.

## Methodology

### Sampling of rhizosphere soil

To identify whether the outbreak of GTDs might be associated with shifts in the composition of rhizosphere microbes, ten vineyards were sampled in August 2022 along the German side of the Upper Rhine representing Northern (Rauenberg), central (Ringsheim), and Southern (Eichstetten, Ihringen) domains within the viticulture region Baden. The majority (eight sites) comprised the commercially important variety Müller-Thurgau, and two sites the traditional variety Silvaner (**Fig. 1**). Rhizosphere soil was collected at 20 cm below the surface from the root-hair zone of plants that either displayed GTD symptoms or were asymptomatic. The soil was immediately transferred to dry ice and remained there during transport, before long-term storage at -80°C. Each vineyard is represented by six rhizosphere samples, three samples from symptomatic, and three from asymptomatic plants.

**Fig. 1:**
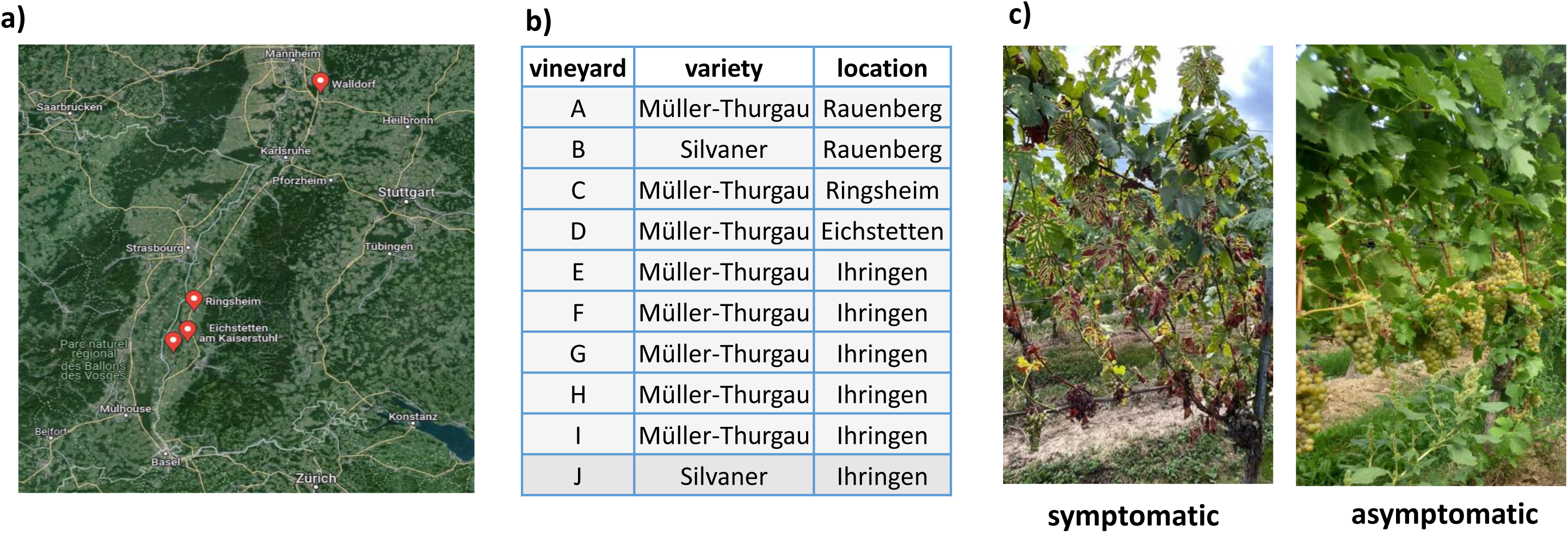
Geographic location (**a**) and grapevine varieties (**b**) in the sampling population of ten representative vineyards from four different geographical locations of the Upper Rhine Valley, Southwest Germany. **c**) representative figures for the asymptomatic and symptomatic vines observed in each vineyard.

### Extraction of DNA and amplicon sequencing

Soil DNA was extracted from aliquots of 400 mg soil using the DNeasy PowerSoil Pro Kit (Qiagen, Hilden-Germany) following the instructions of the manufacturer. Phenolic compounds were removed by washing the DNA with 10% v/v of sodium acetate, then, DNA concentration was quantified using the Qubit® 3.0 fluorometer (Thermo Fisher Scientific) with the Qubit™ dsDNA HS Assay Kit, and quality assessed spectrophotometrically (NanoDrop™ 2000/2000c spectrophotometer, Thermo Fisher Scientific). To analyse the taxonomic structure of the microbial community, 10 ng of the purified DNA were used as template to either amplify 16S ribosomal RNA gene of prokaryotes, or the Internal Transcribed Spacer (ITS) of fungi. For the 16S rRNA, V4-V5 region was targeted using, 0.16 µM of the primer set 5’-GTGCCAGCMGCCGCGGTAA-3’ and 5’-CCGTCAATTCCTTTGAGTTT-3’ ligated with the Illumina adapter. For the ITS, the ITS2 region was addressed with the same concentrations of primers 5’-GCATCGATGAAGAACGCAGC-3’ and 5’-TCCTCCGCTTATTGATATGC-3’. To increase specificity, the PCR was conducted using touchdown cycling at 52-56°C. Annealing took place at 55°C. After PCR, an amplification step was included using 0.04 U/µL of Q5 High-Fidelity DNA Polymerase in presence of Q5 High GC Enhancer (Thermo Fisher). Amplicons were then cleaned up with the DNA Clean & Concentrator Kit (Zymo research, Germany) and subsequently used to prepare amplicon sequencing libraries. Here, 110 ng of cleaned amplicon were selected for a fragment size of 400-600 bp in two steps using 0.4x and 0.7x Agencourt Ampure XP beads (Beckman Coulter). Upon size selection, amplicons were ligated to dual index primers NEBNext® Multiplex Oligos for Illumina® (New England Biolabs, Frankfurt, Germany) following the protocol of the manufacturer, and cleaned afterwards using Ampure XP beads. The prepared libraries were diluted, pooled for equimolarity, and sequenced on a Illumina Novaseq platform to generate 150000 pair-end reads per sample (2 × 250 bp) (Novogene, Munich, Germany).

### Soil chemical analysis

For every vineyard, 6 soil cores were pooled to form a composite representative sample. Soil samples were then sent for chemical analysis using standards assays on soil type, pH, as well as content of macronutrients and micronutrients (Agricultural Analytical and Research Authority of the State of Rheinland-Pfalz, Speyer) following the rules of the German Fertiliser Regulation (DüV).

### Analysis of sequence reads

The obtained paired-end reads from the Illumina sequencer were subjected to quality assessment using the FASTQC tool (Andrews, 2010). Low-quality reads and Illumina adapters were trimmed, and subsequently merged using FASTP (Chen *et al*., 2018). The merged fastq reads were further denoised to filter out chimeric reads as well as reads shorter than 250 bp using the DADA2 plugin in the QIIME2 pipeline based on the denoise-single method (Bolyen et al., 2019). The samples were then mapped to the corresponding sequences and their frequencies calculated, and the resulting operational taxonomic units (OTUs) were then classified either using the database Silva_99 (Robeson et al., 2021) for 16S reads, or the UNITE database (Abarenkov et al., 2024) for reads of fungal ITS. The fungal OTUs were classified with respect to their trophic mode using the FUNGuild database (Nguyen et al., 2016). Since fungal taxa associated with GTDs are not specified in this or alternative databases, we classified, for the current study, wood-trophic taxa that had been previously reported as GTD causal agents (Li et al., 2023a; Martín et al., 2022) as GTDs community. To visualise the high complexity, heatmaps were constructed based on relative frequencies using the ComplexHeatmap software and the circlize tool, clustering variants of rows and columns variants based on their Euclidean distances (Gu, 2022).

### Statistical analysis

To identify rhizomicrobiome members correlated with the outbreak of GTDs, we probed for potential differential abundance among the Müller-Thurgau vineyards using the Analysis of Compositions of Microbiomes with Bias Correction (ANCOM-BC) tool implemented in R. This tool has been developed to derive statistically consistent parameters on the base of samples that differ in size (Lin & Peddada, 2020). Here, the status of the plant (asymptomatic versus asymptomatic) was set as covariate of interest, while the OTUs classified with respect to their role in GTDs were scored per rhizosphere sample. Significantly shifted OTUs were then plotted on a log-linear scale over the plant status to yield log-fold changes, test statistics, standard errors, *P* values, adjusted *P* values, and differential abundance.

### Diversity metrics and correlations analyses

To quantify differences in the composition of the rhizomicrobiome in relation to chemical soil profiles and GTD outbreak, we used several parameters. To address α-diversity (the diversity in a given location), we used the Shannon index as overall estimate (Shannon, 1948). To account for the fact that rare OTUs might be underrepresented due to sampling bias, we also calculated the Chao1 indices (Chao, 1987), and the Faith Phylogenetic Diversity (Faith_PD) index, a parameter that also considers phylogenetic relationships between the taxa (Faith, 1992). All these parameters were calculated using the respective QIIME2 tools. The non-parametrical Kruskal-Wallis test was used to test statistical significance for differences in the a-diversity indices over chemical profile of the soil and the GTD symptomatics.

As alternative approach to assess β-diversity (i.e., differences between different locations), we conducted a Principal Coordinate Analysis to detect commonalities between the sites. For parametrisation, we used here either the Bray-Curtis distance by means of the We used the *vegan* package for R for ecological analyses (Oksanen et al., 2024)., or the Weighted-Unifrac distance (implemented in QIIME2). For visualisation, we employed differentially coloured polygons through the ggplot2 plugin of R, and the stat_ellipse command set at a confidence level of 95%. To classify the chemical profiles of the vineyards, a Principal Component Analysis was carried out using the FactoMinor and factoextra packages of R. In addition, Pearson correlations between rhizomicrobiome diversity metrics and soil chemical profile were calculated and plotted using Hmisc and corrplot packages of R (Harrell, 2024; Wei & Simko, 2021).

### Construction of Co-occurrence networks

Changes in the rhizomicrobiome dynamics and interactions under GTDs outbreak were studied by calculating co-occurrence networks either for asymptomatic or symptomatic vines with a resolution to the genus level. Correlation networks were assessed and visualized using R packages phyloseq (McMurdie & Holmes, 2013), microbiome (Lahti & Shetty, 2017), Hmisc (Harrell, 2024), igraph (Csardi & Nepusz, 2006), and ggplot2 (Wickham, 2016). Fungal and prokaryotic community networks were constructed at the genus level. After elimination of non-annotated OTUs, pairwise correlations were determined using Spearman’s correlation coefficients, filtered based on thresholds of *P* < 0.05 for statistical significance and |r| > 0.6 for correlation strength.

## Results

### Differences in chemical profile are reflected in differences of the rhizomicrobiome

To detect potential shifts in the rhizomicrobiome depending on the chemical profiles of the soil, we probed the rhizosphere of grapevines in ten vineyards along the German side of the Upper Rhine representing the Northern, the central, and the Southern part of the viticulture region Baden (**Fig. 1**). A Principal Component Analysis of soil chemical properties (**Suppl.Fig. 1a**) revealed several types of chemical profile. Soils of vineyards C, D, I, and J showed similar chemical characteristics. Vineyards G and H were categorized separately, mostly due to their low levels of organic Carbon (C), Nitrogen (N), and Boron (B), whereas vineyard F exhibited the opposite profile, characterized by elevated concentrations of these elements. In addition, vineyards E and B were clustered together with comparably high contents of Mn and K (**Suppl.Fig. 1; Suppl.Fig. 2**).

To study the rhizomicrobiome structure in the context of such diverse soil characteristics, amplicon sequencing analysis was performed, using the 16S rRNA for prokaryotes, and the Internal Transcribed Spacer (ITS) for fungi. Following the removal of low-quality and chimeric reads, a total of 8,345,161 fungal ITS reads were processed from 60 samples, identifying 8,893 featured fungal operational taxonomic units (OTUs). On the phylum level, the taxonomic structure was relatively comparable between the vineyards. The most dominant fungi were the *Ascomycota* (between 73% in vineyard D, up to 93% in vineyard G), followed by *Basidiomycota*. Furthermore, the phylum *Rozellomycota* was more prevalent in Vineyard F (**Suppl.Fig. 1b**). Among the twenty most dominant fungal genera, seventeen belonged to the phylum Ascomycota. In addition, *Fusarium* was the most abundant across all vineyards, shaping 12–23% of the total fungal community. Based on the most dominant fungi, the Rauenberg vineyards clustered together using Euclidean distance (**Fig. 2a**), mirroring the pattern observed in the PCA of soil chemical properties (**Suppl.Fig. 1a**). These two vineyards exhibited the lowest relative abundance of Fusarium, but higher levels of two other pathogenic genera, *Penicillium* and *Dactylonectria*, as well as an elevated presence of the beneficial fungus *Solicocczyma*, known to promote root growth (Albornoz et al., 2025). The Eichstetten vineyard also showed distinct profiles of other two fungi, *Fusidium* and *Subulicystidium* (**Fig. 2a**).

**Fig. 2:**
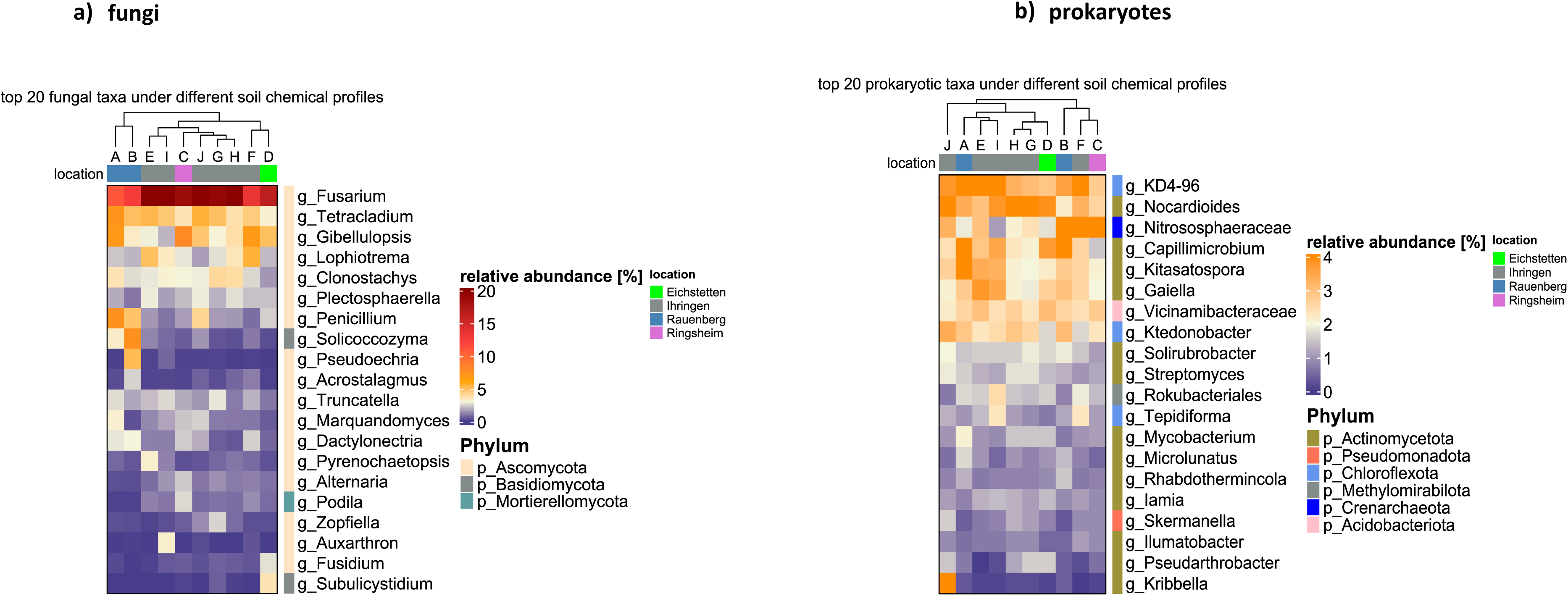
Taxonomic composition of dominant microbial taxa in vineyards rhizomicrobiome across the Upper Rhine Valley; **a**) Relative abundance of the top 20 fungal taxa based on ITS rDNA sequences. **b**) Relative abundance of the top 20 prokaryotic taxa based on 16S rRNA gene sequences. Rows are color-coded according to the taxonomic phylum of each OTU, while columns are color-coded to indicate the geographical origin of each vineyard.

The 16S amplicons reads exhibited higher chimeric read rates, with approximately 30% of reads filtered out. Following denoising step, 3,796,969 high-quality 16S reads remained, representing 84,221 prokaryotic OTUs. Here, the prokaryotic community in the vineyard rhizosphere was significantly enriched, with 10 times more OTUs than observed in the fungal community. At the species level, 1566 Amplicon Sequence Variants (ASVs) of the fungal community were identified, along with 2786 ASVs of prokaryotic origin. Among the prokaryotes, Actinomycetota were the dominant bacterial phylum shaping the rhizobacteriome across the vineyards, with relative abundances ranging from 36% in vineyard I up to 51% in in one vineyard in Ihringen (J), despite similarity of the two vineyards with respect to soil characteristics (**Suppl.Fig. 1a**). *Pseudomonadota, Chloroflexota*, and *Acidobacteriota* were next in relative abundance, respectively (**Suppl.Fig. 1c**). In terms of archaeal communities (overall constituting only a minor fraction), the *Crenarchaeota* were more abundant in vineyards B and F. Additionally, analysis of the twenty most dominant bacterial genera in the vineyard rhizomicrobiome showed that fourteen belonged to the phylum *Actinomycetota*. Despite this, the most dominant genus overall was *KD4*-*96* from the phylum *Chloroflexota*, followed by *Nocardioides* (**Fig. 2b**). Notably, the two vineyards from Ihringen (I and F) exhibited distinct profiles characterized by higher abundances of *Tepidiforma* and *Rokubacteriales*, while vineyard (J) in particular showed a pronounced enrichment of *Kribbella*.

### Wood-colonising fungi dominate in the vineyard rhizomicrobiome

To evaluate the ecological impact of the fungal taxa in the rhizomicrobiome and their potential to colonize woody tissues of grapevine, the defined OTUs were annotated resolving to the genus level using the FUNGuild database, which classifies fungal taxa based on their trophic mode (saprotroph, symbiotroph, or pathotroph), their associated hosts, and, in case of pathogens, their preferred colonization target (wood, root or leaf). Around half of the fungal taxa were pathotrophs (**Fig. 3a**). Most of them were opportunistic pathotrophs, otherwise living as saprotrophs or symbiotrophs, suggesting that their function might vary depending on the condition of the host. In contrast, beneficial taxa (non-pathogenic with symbiotic potential) were found to be less prevalent compared to pathogenic taxa, ranging from only 2.9% to 7.4% (**Fig. 3a**). As alternative approach, we investigated the relative abundance of wood-trophic taxa including those classified as GTDs according to Li et al. (2023a) and Martín et al. (2022). The highest (84-94%) incidence of pathogenic wood-colonising fungi was found in the Southern part of the sampling area, in the vineyards of Ihringen, and Ringsheim,(E, I, And C) whereas the vineyards of Rauenberg (A, and B) in the Northern part harboured less wood pathotrophs, comprising 68-74% of the wood trophic community. By contrast, the Rauenberg vineyards showed the highest abundance of the non-pathogenic saprotrophs (**Fig. 3b**).

**Fig. 3:**
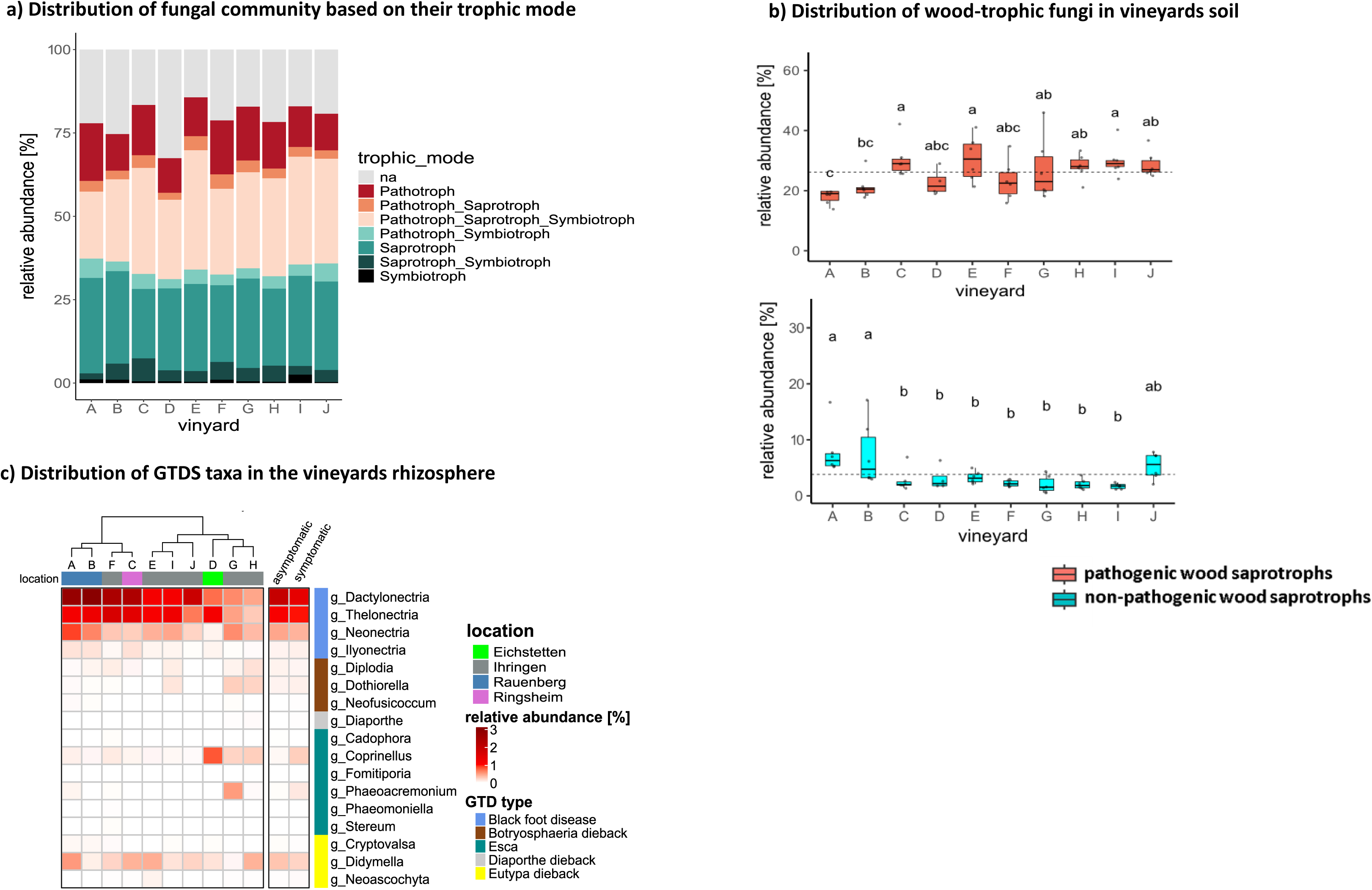
Fungal community annotation in vineyard rhizomicrobiomes across varied geographical and soil chemical profiles. **a**) Annotation of ITS sequences by trophic mode using FUNGuild database. **b**) Relative abundance of wood-trophic OTUs categorized as pathogenic (coral) or non-pathogenic (cyan) to the plants. Significant differences denoted by different letters (Duncan test, P > 0.05). **c**) Heatmap displaying relative abundance of seventeen GTD-annotated OTUs per vineyard or vine status (asymptomatic vs. symptomatic), irrespective of cultivar or location. GTD taxa are color-coded according to associated GTD type.

### Abundance of GTD-associated fungi depends on vineyards, but not on symptom expression

To test, how the abundance of GTD-associated fungi in the rhizosphere relates to the respective vineyard and to the expression of GTD symptoms, we focussed on OTUs from the ITS amplicon sequencing that have been annotated as associated with GTDs (Li et al., 2023a; Martín et al., 2022). In fact, seventeen taxa could be detected in the vineyard rhizosphere linked with GTDs of different type, including Black Foot Disease, Botryosphaeria Dieback, ESCA, Eutypa Dieback, or Diaporthe Dieback. Here, the taxa associated with Black Foot Disease, such as g_*Dactylonectria*, *g_Thelonectria*, and *g_Neonectria* were generally the most prevalent GTD. They were more abundant in the vineyards of Rauenberg (A and B), in the Northern part of the transsect that shared a similar overall profile of GTD-associated fungi, reported by their clustering in terms of Euclidean distance (**Fig. 3c**). Contrasting with Black Foot Disease, fungi associated with Botryosphaeria Dieback (*g_Diplodia, g_Dothiorella*, and *g_Neofusicoccum*) were significantly rarer, and found mainly in vineyards G, H, and I, near Ihringen in the Southern part of the transsect, characterised by *loess* soils and a warm and dry climate. The third disease, Esca, was represented by six OTUs: *g_Coprinellus*, *g_Cadophora*, *g_Phaeoacremonium, g_Stereum*, *g_Fomitiporia,* and *g_Phaeomaniella*. Among them, *g_Coprinellus* was the most abundant Esca taxon in all tested vineyards, particularly in vineyards D, G, and H, followed by *g_Phaeoacremonium*, especially in vineyard G. Three OTUs associated with Eutypa Dieback: *g_Cryptovalsa, g_Neoascochyta*, as well as *g_Didymella* which was significantly detected in all vineyards. To a low extent, OTU *g_Diaporthe*, associated with Diaporthe Dieback was found, without a particular vineyard preference. To assess whether the outbreak of GTDs is linked with a higher abundance of GTD taxa, their relative abundance in the rhizosphere of symptomatic versus asymptomatic vines was calculated, pooling over all vineyards. Here, g_*Coprinellus* was the only OTU that showed significant accumulation for symptomatic plants (*P* < 0.01, Kruskal-Wallis test) (**Fig. 3c**). Thus, with exception of g_*Coprinellus*, we do not see any link between disease outbreak and abundance of GTD-associated fungi.

### GTD outbreak correlates with rhizomicrobiome shifts

To test whether GTD outbreak correlated with significant shifts of the rhizomicrobiome, a differential abundance analysis was carried out, using ANCOM-BC. Since the covariate of interest was the health status of the vine, all rhizosphere samples were pooled into two categories, symptomatic versus asymptomatic vines. Then, for the shifted OTUs, the Log Fold Changes (LFC) for symptomatic over asymptomatic vines, and their statistical significance were calculated, regardless of the cultivar or geographical location. This approach revealed significant shifts of both, the fungal (**Fig. 4a**), and the prokaryotic (**Fig. 4b**) rhizomicrobiome that were also dependent on the chemical properties of the soil. These shifts are described in the following:

**Fig. 4:**
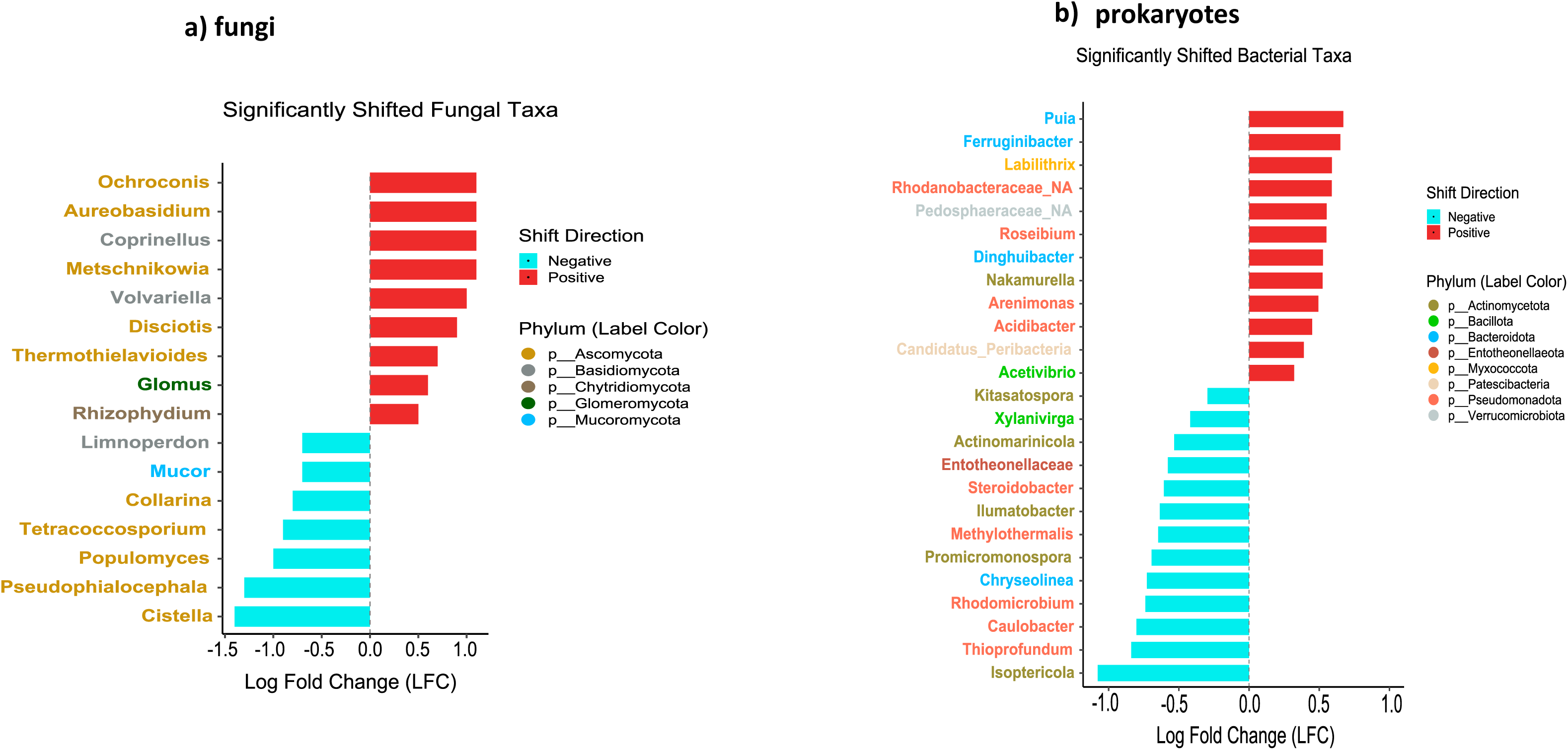
Differentially abundant microbial taxa in vineyards rhizomicrobiome associated with grapevine trunk disease (GTD) outbreak. **a**) Fungal and **b**) bacterial taxa exhibiting significant abundance differences between symptomatic and asymptomatic vines as determined by ANCOM-BC analysis. Log-fold change (LFC) values indicate the magnitude of differential abundance; positive LFC values refer to taxa enriched in symptomatic vines relative to asymptomatic vines. Taxa labels are color-coded by their respective phyla, highlighting shifts in community composition potentially linked to GTD outbreak.

### Fungal shifts

In the fungal community, seven OTUs at the genus level were depleted in symptomatic vines (**Fig. 4a**). Five of these belong to the *Ascomycota*: *Cistella*, *Pseudophialocephala*, *Populomyces*, *Tetracoccosporium*, and *Collarina*. Additionally, two genera from different phyla shifted in parallel: *Mucor* from *Mucoromycota*, and *Limnoperdon* from *Basidiomycota*. On the other hand, nine OTUs were enriched under disease outbreak; Among them were *g_Coprinellus*, proposed as driver of Esca, as well as the epiphytic pathogenic fungus, *g_Aureobasidium*.

To assess how soil chemical profiles correlate with the abundance of rhizomicrobiome fungi associated to the asymptomatic phase, we calculated Pearson correlations between these taxa and individual soil nutrients, adding those fungal phyla that had been found to be generally abundant in the rhizomicrobiome (**Fig. 4a**). We saw significant associations of specific fungal taxa with specific traits of soil chemistry (**Fig. 5a**). For instance, the phylum *Basidiomycota* was positively correlated with the micronutrients Fe, Cu, and Zn, but negatively correlated with CaCOL and pH (alkalinity). Likewise, the phylum *Rozellomycota* showed positive correlations with Org_C and N, but also with the micronutrient Boron (B). Generally, the taxa that decreased during disease outbreaks, seemed more responsive to soil micronutrients.

**Fig. 5:**
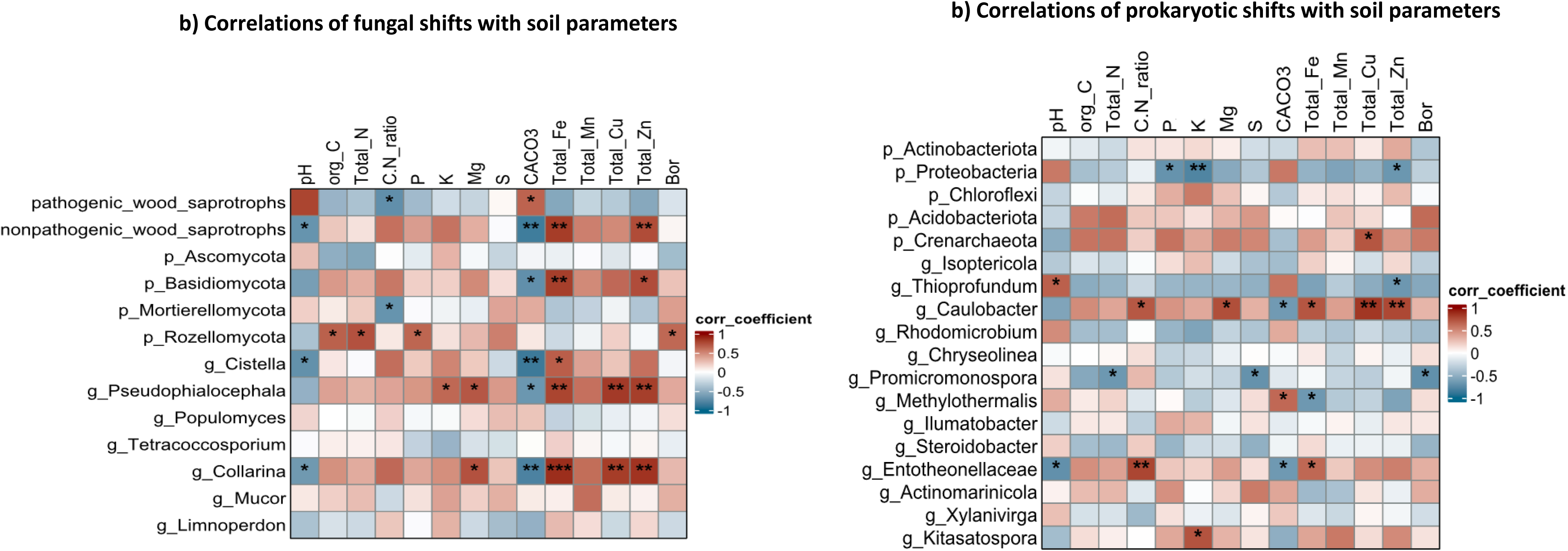
Correlations between rhizomicrobiome composition and soil physicochemical parameters. **a**) Pearson correlation patterns between wood saprotrophs, dominant fungal phyla, and differentially abundant fungal taxa (between symptomatic and asymptomatic vines) with measured soil parameters. **b**) Pearson correlation patterns of dominant bacterial phyla and taxa negatively associated with symptomatic vines in relation to soil variables. Asterisks *, ** and *** refer to significant levels P > 0.05, 0.01, and 0.001, respectively.

Here, *Collarina* as the taxon with the largest fluctuations with respect to soil properties exhibited positive correlations with Zn, Cu, Mn, Fe, Mg, and C.N ratio, followed by g_*Pseudophialocephala*, with significant correlations with Zn, Cu, Fe, Mg, and K. Also, for *Cistella* a link with Fe levels was observed. It is worth noting that fungal members that displayed positive correlations with Fe, became scarce for increases in CaCO_3_ and pH. **Prokaryotic shifts**: The abundance of many bacterial OTUs dropped significantly in symptomatic vines (**Fig. 4b**). Among those taxa that correlated with healthy vines, the two phyla *Actinomycetota* and *Pseudomonadota* exhibited five OTUs that were depleted in symptomatic plants. The most affected taxa were *Isoptericola*, *Thioprofundum*, *Caulobacter*, *Rhodomicrobium,* as well as *Chryseolinea* from *p_Bacteroidota*. Other phyla had only one depleted OTU. For instance, in *p_Bacillota* only one OTU (*Xylanivirga*) decreased, as well as in *p_Entotheonellaeota* (*Entotheonellaceae*). On the other hand, there were also several rhizobacteriome members which accumulated significantly in symptomatic vines: *Puia*, and *Ferruginibacter*, from *p_Bacteroidota*, were enriched most, but also four OTUs from *Pseudomonadota*, as well as a single OTU from each of the phyla *Verrucomicrobiota, p_Patescibacteria* and *Myxococcota* were significantly increased.

We searched for chemical properties of the soil that correlated with these changes of bacterial abundance, but also with dominance of specific prokaryotic phyla independent of disease symptomatics **(Fig. 5b)**. The generally most dominant bacterial phylum, *p_Actinomycetota*, exhibited a negative correlation with only CaCOL while *p_Pseudomonadota* were negatively correlated with P, K, and Zn. Positive correlations were seen for the p_*Acidobacteriota* with B and for *p_Crenarchaeota* with Cu. Soil properties had also a significant impact on several taxa associated with the asymptomatic phase of GTDs. Specifically, *g_Caulobacter* displayed a positive correlation with C.N ratio, as well as with Mg, Fe, Cu, and Zn, but a negative correlation with CaCO_3_ **(Fig. 5b)**. Along with *Caulobacter*, also *Entotheonellaceae* became enriched depending on C.N ratio, Fe, and CaCO_3_, but were depleted in alkaline pH. By contrast, *Thioprofundum*, an OTU from the *Pseudomonadota*, was the only taxon exhibiting a positive correlation with pH.

### Rhizomicrobial co-occurrence networks shift depending on GTD symptoms

To assess whether the interactions and relationships among different rhizomicrobiome taxa are influenced by the health status of the vine, co-occurrence networks were inferred for both, the fungal (**Fig. 6a,b**) and the prokaryotic (**Fig. 6d,e**) microbiome in symptomatic versus asymptomatic vines. Here, the shift of the co-occurrence networks responded qualitatively different in fungi versus prokaryotes. While 1680 significant correlations were detected among the fungal taxa in healthy vines, there was an increase to 1856 significant correlations under GTD outbreak (**Fig. 6g**). A salient component of this increase was the doubling for correlations of GTD-associated taxa with other fungal taxa upon host transition to the symptomatic phase (**Suppl_table1; Fig. 6c**). Here, most GTD taxa sharply changed their correlation profiles. For instance, *Fomitiporia* displayed thirteen significant correlations under disease outbreak, but none in asymptomatic plants. Likewise, correlations of *Stereum* were amplified 8-fold. Furthermore, this fungus extended its associations with other GTD taxa, such as *Phaeomoniella* and *Fomitiporia*, as well as with other six wood-saprotrophic and pathogenic taxa **( Suppl_table1)**. Likewise, *Phaeomoniella*, in symptomatic vines, displayed 29 correlations, not only with *Fomitiporia*, but also with eleven other pathogenic taxa, contrasting with only 10 correlations in healthy vines. The aggressive genus, *Neofusicoccum,* responsible for Botryosphaeria dieback, entertained 15 different correlations during GTD outbreak, compared to only 8 in the asymptomatic phase. For Diaporthe dieback linked with *g_Diaporthe*, a significant correlation, with *g_Paurocotylis*, was only seen in symptomatic vines (**Suppl_table1**). The inverse case, where associations between two pathogens turned loose during GTD outbreak, was far rarer – here, the significant correlation between *Phaoeoacremonium* and *Stereum* was detected only in healthy vines. A third case, where the pathogenic partner of a GTD fungus was swapped by another pathogenic partner, is represented by *Coprinellus*, which was more prevalent in symptomatic vines (**Fig. 4a**). This fungus correlates with *g_Keissleriella* in asymptomatic vines, but switches to a significant correlation with *g_Tulasnella* under the conditions of a GTD outbreak.

**Fig. 6:**
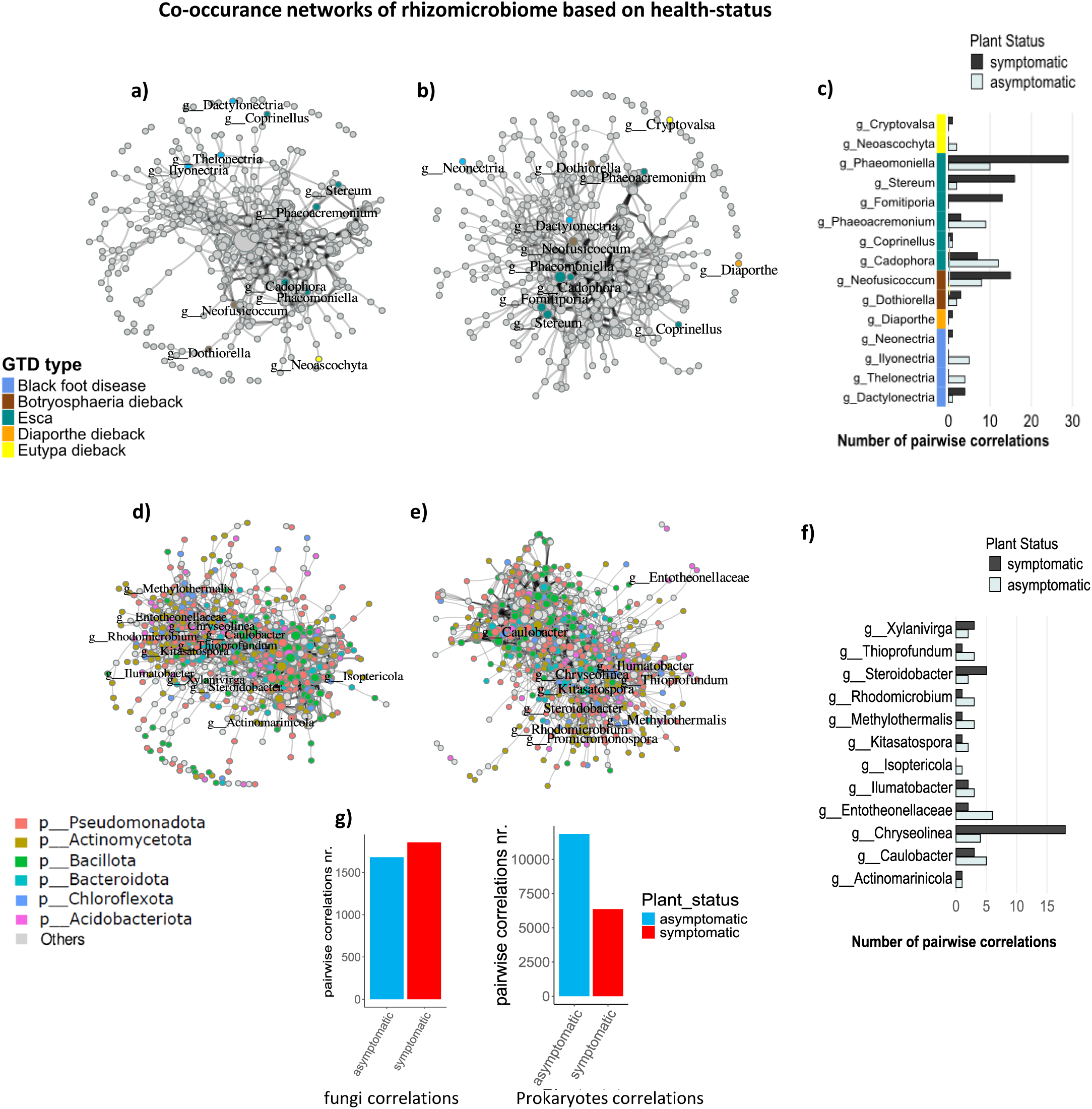
Shifts in rhizomicrobiome co-occurrence networks in vineyards in relation to vine health status: Co-occurrence networks were constructed separately for fungal (**a, b**) and prokaryotic (**d, e**) communities associated with asymptomatic (**a, d**) and GTD-affected (**b, e**) vines. Networks were inferred using Spearman correlations, filtered for statistical significance (p<0.05) and correlation strength (|r|>0.6). Each node represents an operational taxonomic unit (OTU), the size of the node represents the number of the positive pairwise correlations, and each edge represents a significant positive correlation between OTUs. In the fungal networks, GTD-associated taxa are color-coded by disease type, while non-GTD taxa are shown in gray. Prokaryotic nodes are color-coded according to the six most dominant bacterial phyla. OTUs labeled in the prokaryotic networks correspond to taxa significantly reduced under GTD outbreak. **c**) Number of significant pairwise correlations for GTD fungal taxa in the rhizosphere of asymptomatic vs. symptomatic vines, with color codes reflecting GTD types. **f**) Number of significant pairwise correlations for negatively shifted bacterial taxa of rhizomicrobiome under GTD outbreak in the rhizosphere of asymptomatic vs. symptomatic vines. **g**) Total number of significant pairwise correlations within fungal and prokaryotic communities in the vineyards rhizosphere based on the health status.

Contrasting with fungi, the connectivity for the 1167 prokaryotic genera, was drastically decreased from 11856 significant correlations in the healthy vines to 6361 significant correlations under GTDs outbreak **(Fig. 6g; Suppl_table2)**. Here, taxa associated with the asymptomatic phase showed different interaction profiles with other soil prokaryotes. For instance, *Entotheonellaceae* constituted a core node with six strong pairwise correlations in the rhizosphere of healthy vines, but upon GTD outbreak turned into a peripheral node with only two correlations. Also, *Isoptericola* and lost their correlations and even disappeared from the co-occurrence network, while *Thioprofundum, Kitasatospora, Rhodomirobium, Methylothermalis, Illumatobacter, and Caulobacter* robustly lost their positive correlations with other bacteria in symptomatic plants (**Fig. 6f; Suppl_table2**). Only few taxa showed an inverse pattern: *Steroidobacter and Chryseolinea* established more positive correlations only in symptomatic vines.

### Bacterial diversity depends mainly on soil, fungal biodiversity also on geography

As markers for ecosystem robustness, we determined a panel of diversity metrics for the rhizomicrobiome over the different vineyards with their differences in soil parameters and geographical location, either in asymptomatic plants or under disease outbreak. We did not observe any salient changes in the diversity metrics, neither of prokaryotes, nor of fungi in association with GTD outbreak (**Suppl.Fig. 2**). Only for two vineyards, E and G, Shannon entropy, an alpha-diversity index, which accounts for the species richness and evenness within an ecosystem (here, environmental sample), shifted significantly in the bacterial community. Bray_Curtis distance as a quantitative index for community diversity between samples, did not reveal any significant differences in bacteria community between asymptomatic symptomatic vines, but it was significantly shifted in the rhizosphere fungi of symptomatic vines in two other vineyards, D and F.

To link rhizomicrobiome richness with soil parameters, alpha diversity was characterised using Shannon, Chao1, and Faith’s Phylogenetic Diversity (PD) indices over the different vineyards (**Suppl.Fig. 3**), and then correlated either with macronutrients (**Suppl.Fig. 4**) or micronutrients (**Fig. 7**). Generally, the Shannon index remained relatively stable across vineyards, with exceptions observed in the fungal community of vineyard I, and the bacterial communities of vineyards C and E. This index showed no significant correlation, neither with macronutrient nor micronutrient levels. Similarly, Chao1, a richness index that estimates diversity based on abundant taxa, revealed no significant correlation with soil properties, although the prokaryotes in vineyards C and E were significantly different from the other vineyards. Thus, parameters estimating diversity based merely on differences in abundance of taxa, not considering their identity, remained inconspicuous. The outcome changed, when also phylogenetic relationships were included into the parametrisation. Here, Faith’s PD, which considers the phylogenetic differences among taxa, was the most variable parameter among the alpha diversity metrics for both, fungal and prokaryotic, communities (**Suppl.Fig. 3**). This was especially pronounced in the prokaryotic community, where Faith’s PD showed a positive correlation with the principal component of soil properties, Fe, Cu, Zn (**Fig. 7**), and Mg (**Suppl.Fig. 4**).

**Fig. 7:**
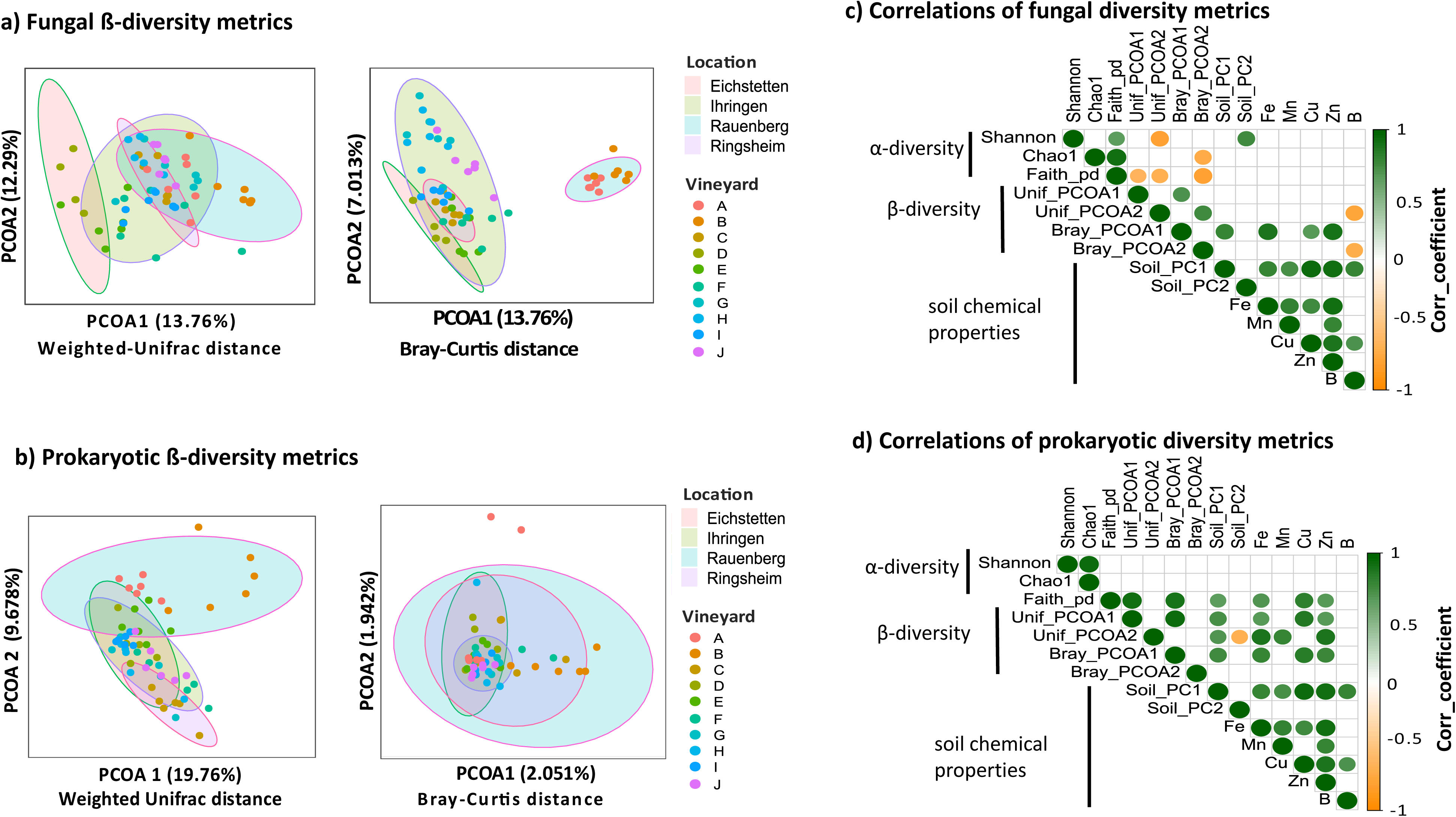
Changes in beta-diversity metrics of the fungal (a) and prokaryotic (b) communities under different geographical locations and soil chemical properties. Principle coordinate analysis (PCOA1, PCOA2) based on either Weighted-Unifrac distance or Bray-Curtis distance for the fungal and prokaryotic communities of different vineyards soils repsectively. β-diversity metrics per geographical location (Eichstetten, Ihringen, Ringsheim, Walldorf) were separated into different colored polygons. To statically evaluate the significance of distant polygons “β-diversity metrics per location”, the stat_ellipse function from the ggplot package was set to a level of 0.95. Pearson correlations of the diversity metrics for the fungal **(c)** and prokaryotic **(d)** communities were calculated in relation to the changes in soil chemical paramters, such as the principal component analysis of soil chemical properties (PC1, PC2) and the micronutrients concentrations. The colorcode of the correlation plots represents the value of the correlation coefficient with significance level= 0.05; green color refers to significantly positive correlation coefficient; orange color represents significantly negative correlation, while the blank color represents the insignificant values of correlation coefficient.

Beta diversity, which evaluates the dissimilarities among different ecosystems, was characterized through Principal Coordinate Analysis (PCoA). Again, this analysis was either conducted either disregarding phylogenetic relationships, based on the Bray-Curtis distance (**Fig. 7b**), or incorporating taxonomic distance, using Weighted-Unifrac distance (**Fig. 7a**). As already seen for alpha diversity, the integration of phylogenetic distance revealed differences that otherwise would have gone unnoticed. While only 4% of prokaryotic beta-diversity could be explained by the first principal PCoAs on the base of Bray-Curtis distance (**Fig. 7b**), the first two Unifrac PCoAs accounted for 30% of the observed diversity (**Fig. 7a**). This improved sensitivity allowed to detect several interesting features: Here, beta-diversity of fungi was clearly different depending on geographical location (**Fig. 7a**). For instance, the selected vineyards in Rauenberg and Ringsheim in the North of the study area were significantly different from those in Eichstetten in the South. However, this difference in fungal diversity seemed to be uncoupled from soil parameters, with exception of CaCO_3_ (**Fig. 7c, Suppl.Fig. S4**). Even disregarding the phylogenetic relationships among OTUs, using the Bray_Curtis distance, the Rauenberg vineyards were significantly divergent with a strong correlation with soil parameters including PC1, Fe, Zn, K and Cu, respectively (**Fig. 7c, Suppl.Fig. S4**). In contrast to fungal biodiversity, the prokaryotic community seemed more dependent on soil parameters. This dependence was so strong that it was not only seen for the Unifrac, but also for the Bray_Curtis distances, where phylogenetic relationship is left aside. Relevant soil parameters were here Fe, Zn, Mg and Cu (**Fig. 7d, Suppl. Fig. S4**).

## Discussion

This study explored the relationship between the outbreak of grapevine trunk diseases (GTDs) and rhizomicrobiome composition and dynamics over ten vineyards, sampled along a North-South transsect in the Upper Rhine valley. The transition from the latent to the necrotrophic phase of the GTD-associated fungi was accompanied by taxonomic shifts in both, the fungal and the prokaryotic, rhizomicrobiome. Using a panel of diversity metrics, we detected correlations between the incidence of prokaryotic taxa associated with the latent, asymptomatic phase and soil properties, whereas for the respective fungal taxa also geographic location was found to be relevant. We identified taxa associated with the latent “asymptomatic” phase and their interactions with rhizosphere flora and the soil properties. The geographical location and soil properties showed variable effects on the composition and diversity metrics of both fungal and bacterial communities. The analysis of correlation networks revealed that GTD-associated fungi increased their mutual association after outbreak of the disease. These findings stimulate several questions in the context of GTD outbreak that will be discussed in the following: Why are the prokaryotes responsive to soil chemistry, while the fungi are not? What are potential mechanisms behind the association of specific prokaryotes with grapevine health? What does the altered fungal correlation network mean for our concept of health and disease?

### Soil ecology: fungi respond by metabolic state, bacteria by proliferation

The rhizomicrobiome plays a central role for maintaining plant health and resilience against environmental challenges (reviewed in Berendsen et al., 2012; Gu et al., 2022). Therefore, insight into the grapevine rhizomicrobiome structure, as well as environmental key factors shaping its richness, diversity, and the incidence of beneficial taxa is relevant for sustainable viticulture. Along this line, two salient features emerge from our analysis:

For the fungal community, it is host status, rather than soil properties, that seems to define disease outbreak. The dominant fungal phyla in the vineyard rhizosphere were *Ascomycota* and *Basidiomycota*, respectively (**Suppl.Fig. 1**), similar to previous studies performed on different cultivars, different geographical and environmental conditions (Bao et al., 2022; Berlanas et al., 2019; Coller et al., 2019; Lailheugue et al., 2024). In addition, there were no significant differences in the incidence of symbiotrophic or pathotrophic taxa among the different vineyards (**Fig. 3a**). However, around 50% of the detected fungal taxa were multi-trophic with a pathotrophic potential. This is consistent with a scenario, where disease outbreak is not defined by increased abundance of GTD associated taxa, but rather by changes in their behaviour. In fact, such a conditional pathogenesis has been demonstrated to link to changes in secondary metabolites, for instance, in the fungal GTD model *Neofusicoccum parvum* (Khattab et al., 2022; Flubacher et al., 2023). While secondary metabolism in fungi is very rich and elaborate, most of the underlying gene clusters are silent, if not activate by specific (mostly unknown) signals (for review see Brakhage and Schroekh, 2011). It is, therefore, not surprising that, for the rhizomicrobiome fungi, differences in soil ecology will be reflected in changes of metabolic state, rather than in changes of abundance.

In contrast to fungi, prokaryotic phyla seemed more influenced by geography and environmental factors. In our study, *Actinomycetota* emerged as the predominant bacterial phylum, followed by *Pseudomonadota*, *Chloroflexota*, and *Acidobacteriota* (**Suppl.Fig. 1c**). With exception of the *Chloroflexota*, these dominating phyla have been found in other vineyard studies, albeit at variable composition. For example, similar profiles of *Actinomycetota* and *Pseudomonadota* were reported across various cultivars in the grape-growing regions of Huailai County, China by Bao et al. (2022). A strong dependence on geography can also been concluded by differences in the dominant phyla of the grapevine rhizosphere among different regions in Europe. While the *Pseudomonadota* was the dominant phylum, followed by *Actinomycetota* in vineyards located in Bordeaux, France (Lailheugue et al., 2024), Italy (Marasco et al., 2018), and Spain (Berlanas et al., 2019), a different study on ten vineyards across four sites in Italy identified *Acidobacteriota* as most abundant phylum (Coller et al., 2019). Such comparisons between different studies have to be taken with care, because timing of sampling might also affect the rhizomicrobiome structure (Berlanas et al., 2019). However, for the data from the current study, seasonal differences can be excluded, because the samples were collected in August, because outbreak of GTDs becomes fully manifest in late summer. Instead, our data lead to the conclusion that the substantial differences in prokaryotic abundance might derive from soil properties. This is supported, for instance, by the negative correlation between *Pseudomonadota* and content of Zn, K and P, or by the strong positive correlation between *Acidobacteriota* and B and nitrogen content (**Fig. 5b**). Globally, bacterial diversity metrics, especially those incorporating phylogenetic relationships such as Faith_PD and Unifrac, showed a significant correlation with soil properties, especially the content of micronutrients Fe, Cu, Mn, and Zn (**Fig.** 7**c****, Suppl.Fig. 4**). Such elements are key players for enzymes regulating microbial proliferation and many biological processes, e.g. Fe for N fixation, Zn and Cu for immunocompetence, and Fe and Mn for respiration. Both elements, Fe and Mn could also serve as electron donors and acceptors, during soil redox reactions of C, N, and S (Dai et al., 2023b; Dubinsky et al., 2010; Whalen et al., 2018). In contrast to fungi that can respond to environmental conditions by releasing previously silent metabolic modules, bacteria need to respond by altering the composition of their consortia that are often coupled by concerted metabolisation of given substrates, giving rise to the concept of Metabolically Cohesive Consortia (for a conceptual review see Pascual-García et al., 2020).

In shorthand, the fungal communities in the rhizomicrobiome respond by adjusting their metabolic state, while the bacteria respond by altering their proliferation.

### Disease outbreak is associated with shifts in the bacterial rhizomicrobiome

Under disease outbreak, we observed strong shifts in the composition of the bacterial rhizomicrobiome (contrasting with the fungi, which will be discussed in the next section), reflected in the co-occurrence networks. The positive correlations, as reported by edges among the network nodes, dropped drastically almost to the half among (**Fig. 6g**). Salient features were the depletion of *Isoptericola*, *Thioprofundum*, *Caulobacter*, *Rhodomicrobium,* as well as *Chryseolinea* (**Fig. 4b**). These taxa might, therefore, used as indicators for grapevine health. In addition, some taxa lost their positive correlations or completely disappeared from the co-occurrence network, e.g. *Isoptericola* (**Fig. 6e; Suppl. Table 2**). Possible reasons might be increased competition for nutrients, reduced functionality, but also the rise of outbreak-associated bacterial communities. Since around 50% of the fungal flora in the rhizomicrobiome have pathotrophic potential (**Fig. 3a**), the depleted or less connected bacterial networks might be a factor driving the transition to fungal pathogenicity.

While the underlying mechanisms remain to be elucidated for grapevine, current findings from other crops suggest that root exudates released from healthy plants promote the establishment of a rhizomicrobiome that promotes nutrient cycling, enhances plant immunity, and maintains soil health (Chen et al., 2024; Du et al., 2024; Wilhelm et al., 2023). Under the severe stress conditions that often herald a GTD outbreak (Khattab et al., 2022), either the absence or the modification of such root exudates might contribute to the disruption of the established microbial networks, such that beneficial microbiota become depleted, while pathogens become promoted, and conditional fungal pathogens alter their lifestyle towards parasitism. Harnessing the established bacterial networks in health-associated rhizomicrobiomes, (**Fig. 6**), might be a strategy for sustainable biocontrol of GTDs. Micronutrients supplements might help in this regard since they showed strong correlations with phylogenetic diversity metrics as well as with higher abundance of health-associated bacteria (**Fig. 5**; **Fig. 7**). Whether such bacteria are actively promoting plant resilience, or whether they are just attracted to plants endowed with resilience cannot be inferred from a correlative study, of course. H owever, these bacteria are at least candidates, and it is worthwhile to probe them individually for a potential activation of plant immunity. This would also be interesting for application, because those, where activation of immunity can be confirmed could be developed into new sustainable biocontrol agents against GTDs. The potential of this approach has already been by studies, where co-inoculation with members of *Actinomycetota* and *Bacillota* mitigated GTD progression (for review see Cobos et al., 2022). This mitigation could be direct, by allelopathic control. For instance, *Streptomyces* either endophytic sp. VV/E1, or two rhizosphere isolates, sp. VV/R1 and sp. VV/R4, significantly inhibited the growth of *Dactylonectria* sp. and *Ilyonectria* sp., fungi involved in black foot disease, as well as of *Phaeomoniella chlamydospora*, and *P. minimum*, associated with the Esca disease (Álvarez-Pérez et al., 2017). In addition to direct growth inhibition, secreted compounds might also act indirectly, by activating host defence. Likewise, In fact, *Bacillus pumilus* (S32) and *Paenibacillus* sp. (S19) were shown to secrete antifungal volatiles, including 1-octen-3-ol and 2,5-dimethyl pyrazine that not only suppressed the Esca fungus *Phaeomoniella chlamydospora* but also upregulated phytoalexin biosynthesis genes of grapevine (Haidar et al., 2016). In a similar manner, *Bacillus subtilis* PTA-271 could simultaneously enhance defence signalling and reduce the growth rate of *Neofusicoccum parvum* in grapevine (Trotel-Aziz et al., 2019).

### GTD outbreak: a matter of fungal ecology rather than pathogen incidence?

Contrasting with other grapevine diseases, such as Downy Mildew (caused by the oomycete *Plasmopara viticola*), or Powdery Mildew (caused by the ascomycete *Erysiphe necator*), the causal chain in GTDs is far from elucidated. Classically, pathogens are identified through demonstrating that they are necessary and sufficient for symptomatics, an approach known as Koch Postulates (Loeffler, 1884). This classical approach fails for GTDs, because symptoms can rarely be attributed to absence and presence of a given fungus. Moreover, attempts to compare symptomatic vines with asymptomatic controls did not yield significant differences in the composition of the mycobiome. In one of the (few) rigorous comparisons, the authors arrive at the provocative conclusion “What if esca disease of grapevine were not a fungal disease?” (Hofstetter et al., 2012), arguing that the presence of certain wood-decaying fungi in both asymptomatic or symptomatic trunks might be due to their saprotrophic lifestyle breaking down tissue that had died for other reasons, such as overpruning or frost damage. If it is not the mere presence of a microbe that leads to pathogenesis, it might be the conditions that render a microbe into a pathogen. The findings of the current study, mainly the shifts in the correlation networks, support such a contextual model of pathogenesis, clearly transcending the Koch Postulates.

On the one hand, most prevalent GTD associated taxa, namely those reported in the context of black foot disease, showed no significant shifts compared to the asymptomatic phase (**Fig. 3c**). The only apparent candidate, *Coprinellus*, seems to be only a hitchhiker and not a driver. In controlled infection studies, it failed to induce foliar symptoms or trigger disease outbreak contrasting with other Esca fungi (Brown et al., 2020). Moreover, unlike many GTDs taxa which are wood-obligate saprotrophs, *Coprinellus* was detected also in grapevine leaves (Cui et al., 2024). Therefore, *Coprinellus* might rather be a saprotroph, potentially feeding on decayed plant material following the outbreak.

Also for the other sixteen GTD associated taxa, abundance changes after outbreak were insignificant. However, what was significant, was the dynamics of interactions which were amplified within the fungal community, prominently in those with pathotrophic potential (**Suppl_Table 1**). Here, some GTD taxa strongly increased their mutual correlations, especially Esca-associated fungi, such as *Phaeomoniella* with *Fomitiporia*, *Stereum* with *Fomitiporia*, and *Stereum* with *Phaeomoniella*. These patterns are consistent with a model, where these fungi initiate mutualistic interactions and benefit each other through co-colonisation or metabolic-cross feeding. Whether the opportunistic shift of these Esca taxa towards pathogenic behaviour is triggered by a breakdown in plant immunity due to loss of beneficial microbes remains to be elucidated by targeted infection experiments in the presence of tailored rhizomicrobiomes.

### Outlook

It has to be kept in mind that the nature of a agroecological study as the current one is descriptive, and the outcome is confined to correlations. Since the sampling had to be in August, when symptoms are clearly manifest, the sampling represents a static snapshot. To infer the causal chain from such snapshots alone is principally not possible. Nevertheless, several candidates, both for pathogenesis, as well as for salutogenesis, could be identified. To integrate the temporal dynamics of pathogenesis, these candidates need to be tested functionally in controlled infection assays to assess, for instance changes in the root secretome under GTDs outbreak as well as direct promoting or inhibiting interactions between microbes, or activation of plant immunity. The ultimate goal will be to develop new biocontrol agents to prevent or even to cure grapevine trunk diseases.

## Funding declaration and acknowledgement

This work was supported by Microbes for future project (M4F), which was funded by Strategy fund of Karlsruhe Institute of Technology.

The authors have no relevant financial or non-financial interests to disclose. All authors read and approved the final manuscript.

## Data availability

**Sequencing rawdata:** NCBI bioproject (Vineyards_rhizomicrobiome), Accession: PRJNA1328367, ID: 1328367 (https://www.ncbi.nlm.nih.gov/bioproject/1328367)

**Bioinformatics**: Github respiratory, “Vineyards_rhizomicrobiome_Upper_Rhine” (https://github.com/Khattab2022/Vineyards_rhizomicrobiome_Upper_Rhine)

## References

Abarenkov, K., Nilsson, R. H., Larsson, K. H., Taylor, A. F. S., May, T. W., Frøslev, T. G., Pawlowska, J., Lindahl, B., Põldmaa, K., Truong, C., Vu, D., Hosoya, T., Niskanen, T., Piirmann, T., Ivanov, F., Zirk, A., Peterson, M., Cheeke, T. E., Ishigami, Y., … Kõljalg, U. (2024). The UNITE database for molecular identificationãnd taxonomic communication of fungiãnd other eukaryotes: sequences, taxaãnd classifications r econsider ed. Nucleic Acids Research, 52(D1), D791–D797. 10.1093/nar/gkad1039

Albornoz, F., Carvajal, M., Catrileo, D., Gebauer, M., & Godoy, L. (2025). Volatile organic compounds produced after exposure of tomato roots to the soil yeast Solicoccozyma terrea modulate root nitrate transporters in tomato. Plant and Soil. 10.1007/s11104-025-07393-8

Álvarez-Pérez, J. M., González-García, S., Cobos, R., Olego, M. Á., Ibañez, A., Díez-Galán, A., Garzón-Jimeno, E., & Coque, J. J. R. (2017). Use of endophytic and rhizosphere actinobacteria from grapevine plants to reduce nursery fungal graft infections that lead to young grapevine decline. Applied and Environmental Microbiology, 83(24). 10.1128/AEM.01564-17

Bao, L., Sun, B., Wei, Y., Xu, N., Zhang, S., Gu, L., & Bai, Z. (2022). Grape Cultivar Features Differentiate the Grape Rhizosphere Microbiota. Plants, 11(9). 10.3390/plants11091111

Berendsen, R. L., Pieterse, C. M. J., & Bakker, P. A. H. M. (2012). The rhizosphere microbiome and plant health. In Trends in Plant Science (Vol. 17, Issue 8, pp. 478–486). 10.1016/j.tplants.2012.04.001

Berlanas, C., Berbegal, M., Elena, G., Laidani, M., Cibriain, J. F., Sagües, A., & Gramaje, D. (2019). The fungal and bacterial rhizosphere microbiome associated with grapevine rootstock genotypes in mature and young vineyards. Frontiers in Microbiology, 10(MAY). 10.3389/fmicb.2019.01142

Bolyen. E; Rideout J.R; Dillon M.R; Bokulich N.A.; Abnet C.C.; Al-Ghalith G.A.; Alexander H.; Alm E.J.; Arumugam M. (2019). Reproducible, interactive, scalable and extensible microbiome data science using QIIME 2. Nature Biotechnology, 37(8), 850–852. 10.1038/s41587-019-0190-3

Brown, A. A., Lawrence, D. P., & Baumgartner, K. (2020). Role of basidiomycete fungi in the grapevine trunk disease esca. Plant Pathology, 69(2), 205–220. 10.1111/ppa.13116

Chao, A. (1987). Estimating the Population Size for Capture-Recapture Data with Unequal Catchability (Vol. 43, Issue 4). https://www.jstor.org/stable/2531532

Chen, Q., Song, Y., An, Y., Lu, Y., & Zhong, G. (2024). Soil Microorganisms: Their Role in Enhancing Crop Nutrition and Health. In Diversity (Vol. 16, Issue 12). Multidisciplinary Digital Publishing Institute (MDPI). 10.3390/d16120734

Cobos, R., Ibañez, A., Diez-Galán, A., Calvo-Peña, C., Ghoreshizadeh, S., & Coque, J. J. R. (2022). The Grapevine Microbiome to the Rescue: Implications for the Biocontrol of Trunk Diseases. In Plants (Vol. 11, Issue 7). MDPI. 10.3390/plants11070840

Coller, E., Cestaro, A., Zanzotti, R., Bertoldi, D., Pindo, M., Larger, S., Albanese, D., Mescalchin, E., & Donati, C. (2019). Microbiome of vineyard soils is shaped by geography and management. Microbiome, 7(1). 10.1186/s40168-019-0758-7

Cui, S., Zhou, L., Fang, Q., Xiao, H., Jin, D., & Liu, Y. (2024). Growth period and variety together drive the succession of phyllosphere microbial communities of grapevine. Science of the Total Environment, 950. 10.1016/j.scitotenv.2024.175334

Dai, Z., Guo, X., Lin, J., Wang, X., He, D., Zeng, R., Meng, J., Luo, J., Delgado-Baquerizo, M., Moreno-Jiménez, E., Brookes, P. C., & Xu, J. (2023a). Metallic micronutrients are associated with the structure and function of the soil microbiome. Nature Communications, 14(1). 10.1038/s41467-023-44182-2

Dai, Z., Guo, X., Lin, J., Wang, X., He, D., Zeng, R., Meng, J., Luo, J., Delgado-Baquerizo, M., Moreno-Jiménez, E., Brookes, P. C., & Xu, J. (2023b). Metallic micronutrients are associated with the structure and function of the soil microbiome. Nature Communications, 14(1). 10.1038/s41467-023-44182-2

De Vries, F. T., Griffiths, R. I., Knight, C. G., Nicolitch, O., & Williams, A. (2020). Harnessing rhizosphere microbiomes for drought-resilient crop production. 10.1126/science.aaz5192

Dubinsky, E. A., Silver, W. L., & Firestone, M. K. (2010). Tropical forest soil microbial communities couple iron and carbon biogeochemistry. Ecology, 91(9), 2604–2612. 10.1890/09-1365.1

Du, Y., Han, X., & Tsuda, K. (2024). Microbiome-mediated plant disease resistance: recent advances and future directions. In Journal of General Plant Pathology. Springer. 10.1007/s10327-024-01204-1

Faith, D. P. (1992). Conservation evaluation and phylogenetic diversity. In Biological Conservation (Vol. 61).

Field, K. J., Pressel, S., Duckett, J. G., Rimington, W. R., & Bidartondo, M. I. (2015). Symbiotic options for the conquest of land. In Trends in Ecology and Evolution (Vol. 30, Issue 8, pp. 477–486). Elsevier Ltd. 10.1016/j.tree.2015.05.007

Flubacher, N., Baltenweck, R., Hugueney, P., Fischer, J., Thines, E., Riemann, M., Nick, P., & Khattab, I. M. (2023). The fungal metabolite 4-hydroxyphenylacetic acid from Neofusicoccum parvum modulates defence responses in grapevine. Plant Cell and Environment, 46(11), 3575–3591. 10.1111/pce.14670

Fotios, B., Sotirios, V., Elena, P., Anastasios, S., Stefanos, T., Danae, G., Georgia, T., Aliki, T., Epaminondas, P., Emmanuel, M., George, K., Kalliope, P. K., & Dimitrios, K. G. (2021). Grapevine wood microbiome analysis identifies key fungal pathogens and potential interactions with the bacterial community implicated in grapevine trunk disease appearance. Environmental Microbiomes, 16(1). 10.1186/s40793-021-00390-1

Gilbert, J. A., Van Der Lelie, D., & Zarraonaindia, I. (2014). Microbial terroir for wine grapes. In Proceedings of the National Academy of Sciences of the United States of America (Vol. 111, Issue 1, pp. 5–6). 10.1073/pnas.1320471110

Gu, S., Wei, Z., Shao, Z., Friman, V. P., Cao, K., Yang, T., Kramer, J., Wang, X., Li, M., Mei, X., Xu, Y., Shen, Q., Kümmerli, R., & Jousset, A. (2020). Competition for iron drives phytopathogen control by natural rhizosphere microbiomes. Nature Microbiology, 5(8), 1002–1010. 10.1038/s41564-020-0719-8

Gu, Y., Banerjee, S., Dini-Andreote, F., Xu, Y., Shen, Q., Jousset, A., & Wei, Z. (2022). Small changes in rhizosphere microbiome composition predict disease outcomes earlier than pathogen density variations. ISME Journal, 16(10), 2448–2456. 10.1038/s41396-022-01290-z

Gu, Z. (2022). Complex heatmap visualization. IMeta, 1(3). 10.1002/imt2.43

Haidar, R., Roudet, J., Bonnard, O., Dufour, M. C., Corio-Costet, M. F., Fert, M., Gautier, T., Deschamps, A., & Fermaud, M. (2016). Screening and modes of action of antagonistic bacteria to control the fungal pathogen Phaeomoniella chlamydospora involved in grapevine trunk diseases. Microbiological Research, 192, 172–184. 10.1016/j.micres.2016.07.003

Hofstetter, V., Buyck, B., Croll, D., Viret, O., Couloux, A., & Gindro, K. (2012). What if esca disease of grapevine were not a fungal disease? Fungal Diversity, 54, 51–67. 10.1007/s13225-012-0171-z

Khattab, I. M., Fischer, J., Kaźmierczak, A., Thines, E., & Nick, P. (2023). Ferulic acid is a putative surrender signal to stimulate programmed cell death in grapevines after infection with Neofusicoccum parvum. Plant Cell and Environment, 46(1), 339–358. 10.1111/pce.14468

Kwak, M. J., Kong, H. G., Choi, K., Kwon, S. K., Song, J. Y., Lee, J., Lee, P. A., Choi, S. Y., Seo, M., Lee, H. J., Jung, E. J., Park, H., Roy, N., Kim, H., Lee, M. M., Rubin, E. M., Lee, S. W., & Kim, J. F. (2018). Rhizosphere microbiome structure alters to enable wilt resistance in tomato. Nature Biotechnology, 36(11), 1100–1116. 10.1038/nbt.4232

Lailheugue, V., Darriaut, R., Tran, J., Morel, M., Marguerit, E., & Lauvergeat, V. (2024). Both the scion and rootstock of grafted grapevines influence the rhizosphere and root endophyte microbiomes, but rootstocks have a greater impact. Environmental Microbiome, 19(1). 10.1186/s40793-024-00566-5

Lee, S. M., Kong, H. G., Song, G. C., & Ryu, C. M. (2021). Disruption of Firmicutes and Actinobacteria abundance in tomato rhizosphere causes the incidence of bacterial wilt disease. ISME Journal, 15(1), 330–347. 10.1038/s41396-020-00785-x

Lin, H., & Peddada, S. Das. (2020). Analysis of compositions of microbiomes with bias correction. Nature Communications, 11(1). 10.1038/s41467-020-17041-7

Li, Y., Li, X., Zhang, W., Zhang, J., Wang, H., Peng, J., Wang, X., & Yan, J. (2023a). Belowground microbiota analysis indicates that Fusarium spp. exacerbate grapevine trunk disease. Environmental Microbiome, 18(1). 10.1186/s40793-023-00490-0

Li, Y., Li, X., Zhang, W., Zhang, J., Wang, H., Peng, J., Wang, X., & Yan, J. (2023b). Belowground microbiota analysis indicates that Fusarium spp. exacerbate grapevine trunk disease. Environmental Microbiome, 18(1). 10.1186/s40793-023-00490-0

Marasco, R., Rolli, E., Fusi, M., Michoud, G., & Daffonchio, D. (2018). Grapevine rootstocks shape underground bacterial microbiome and networking but not potential functionality. Microbiome, 6(1). 10.1186/s40168-017-0391-2

Martín, L., García-García, B., & Alguacil, M. del M. (2022). Interactions of the Fungal Community in the Complex Patho-System of Esca, a Grapevine Trunk Disease. International Journal of Molecular Sciences, 23(23). 10.3390/ijms232314726

Mendes, R., Kruijt, M., Bruijn, I. de, Dekkers, E., van der voort, M., & Schneider, J. H. M. (2011). Deciphering the RhizosphereMicrobiome for Disease-Suppressive Bacteria. Science, 332(6033), 1093–1097. 10.1126/science.1202007

Nguyen, N. H., Song, Z., Bates, S. T., Branco, S., Tedersoo, L., Menke, J., Schilling, J. S., & Kennedy, P. G. (2016). FUNGuild: An open annotation tool for parsing fungal community datasets by ecological guild. Fungal Ecology, 20, 241–248. 10.1016/j.funeco.2015.06.006

Pascual-García, A., Bonhoeffer, S., & Bell, T. (2020). Metabolically cohesive microbial consortia and ecosystem functioning. In Philosophical Transactions of the Royal Society B: Biological Sciences (Vol. 375, Issue 1798). Royal Society Publishing. 10.1098/rstb.2019.0245

Pollard-Flamand, J., Boulé, J., Hart, M., & Úrbez-Torres, J. R. (2022). Biocontrol Activity of Trichoderma Species Isolated from Grapevines in British Columbia against Botryosphaeria Dieback Fungal Pathogens. Journal of Fungi, 8(4). 10.3390/jof8040409

Ren, B., Wang, X., Duan, J., & Ma, J. (2019). Rhizobial tRNA-derived small RNAs are signal molecules regulating plant nodulation. https://www.science.org

Robeson, M. S., O’Rourke, D. R., Kaehler, B. D., Ziemski, M., Dillon, M. R., Foster, J. T., & Bokulich, N. A. (2021). RESCRIPt: Reproducible sequence taxonomy reference database management. PLoS Computational Biology, 17(11). 10.1371/journal.pcbi.1009581

Shannon, C. E. (1948). A Mathematical Theory of Communication. In The Bell System Technical Journal (Issue 3).

Trotel-Aziz, P., Abou-Mansour, E., Courteaux, B., Rabenoelina, F., Clément, C., Fontaine, F., & Aziz, A. (2019). Bacillus subtilis PTA-271 counteracts botryosphaeria dieback in grapevine, triggering immune responses and detoxification of fungal phytotoxins. Frontiers in Plant Science, 24. 10.3389/fpls.2019.00025

Wei, Z., Gu, Y., Friman, V.-P., Kowalchuk, G. A., Xu, Y., Shen, Q., & Jousset, A. (2019). Initial soil microbiome composition and functioning predetermine future plant health. In Sci. Adv (Vol. 5). https://www.science.org

Whalen, E. D., Smith, R. G., Grandy, A. S., & Frey, S. D. (2018). Manganese limitation as a mechanism for reduced decomposition in soils under atmospheric nitrogen deposition. Soil Biology and Biochemistry, 127, 252–263. 10.1016/j.soilbio.2018.09.025

Wilhelm, R. C., Amsili, J. P., Kurtz, K. S. M., van Es, H. M., & Buckley, D. H. (2023). Ecological insights into soil health according to the genomic traits and environment-wide associations of bacteria in agricultural soils. ISME Communications, 3(1). 10.1038/s43705-022-00209-1

Zarraonaindia, I., Owens, S. M., Weisenhorn, P., West, K., Hampton-Marcell, J., Lax, S., Bokulich, N. A., Mills, D. A., Martin, G., Taghavi, S., van der Lelie, D., & Gilbert, J. A. (2015). The soil microbiome influences grapevine-associated microbiota. MBio, 6(2). 10.1128/mBio.02527-14

