## Supplementary_figures for "Health or disease – a question of rhizomicrobial ecology? The case of Grapevine Trunk Disease"

### Slide 1
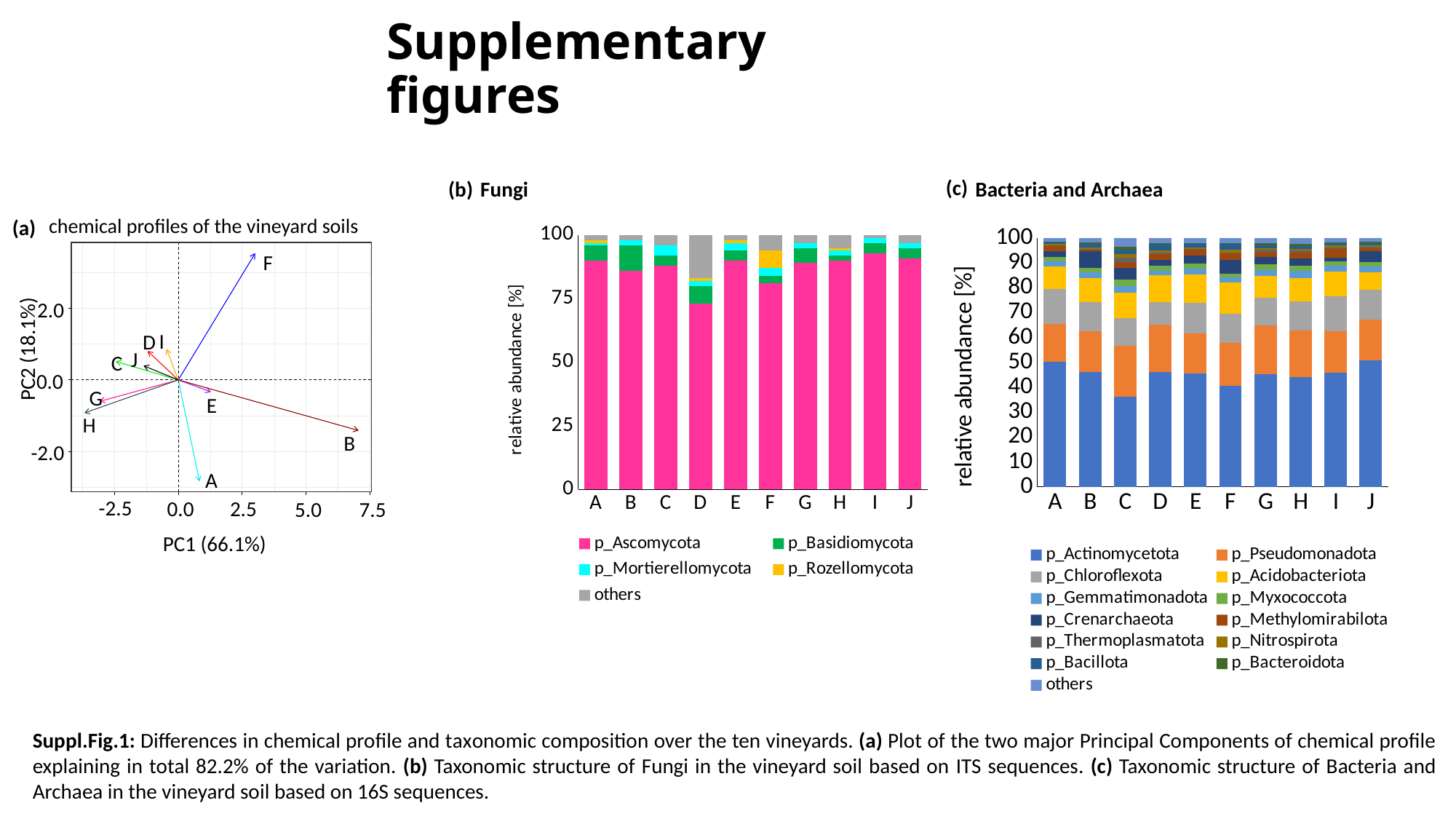

#### Supplementary figures
(c)
(b)
Fungi
Bacteria and Archaea
chemical profiles of the vineyard soils
(a)
###### Chart
| Category | p_Actinomycetota | p_Pseudomonadota | p_Chloroflexota | p_Acidobacteriota | p_Gemmatimonadota | p_Myxococcota | p_Crenarchaeota | p_Methylomirabilota | p_Thermoplasmatota | p_Nitrospirota | p_Bacillota | p_Bacteroidota | others |
|---|---|---|---|---|---|---|---|---|---|---|---|---|---|
| A | 50.082498190062296 | 15.39043094506666 | 13.921734309627816 | 9.187485136054134 | 2.020452634544654 | 1.7636977068719062 | 2.222094967631821 | 2.439836117307223 | 0.0036118033734243507 | 0.6029284536163096 | 0.4081713534210785 | 0.285845136035905 | 1.6712132463867653 |
| B | 46.18772640588113 | 16.183558680345303 | 11.871743227631356 | 9.703505649710955 | 2.2003866094085596 | 1.7210630315830129 | 6.684175959277618 | 1.0081964388134133 | 0.003075443515515877 | 0.5899286102797342 | 1.285714073587352 | 0.7537941073497832 | 1.8071317626162668 |
| C | 36.1556013424011 | 20.496693530839533 | 11.160918540143621 | 10.273290838163748 | 2.4247800240784874 | 2.6331381778203276 | 4.842057119175957 | 2.3010319677515025 | 1.8458433352927246 | 1.3041374924081832 | 1.81727878257087 | 1.19324828002758 | 3.5519805693263455 |
| D | 46.09877521092176 | 18.933253850128647 | 9.225728690484052 | 10.643536609949901 | 1.964487999759486 | 1.9650413063670669 | 2.291518670974323 | 2.4580332340380244 | 0.3514752698220381 | 1.1146719720030562 | 2.173930083823472 | 0.6049067693333976 | 2.174640332394772 |
| E | 45.6411553104682 | 15.896095918739382 | 12.4490886093144 | 11.388152006724482 | 2.3717895478327518 | 1.841298363161919 | 3.2431070498769086 | 2.3572087530794152 | 0.23633790518426856 | 0.7605565808437621 | 1.2482763463124087 | 0.30924993826510433 | 2.2576836701969976 |
| F | 40.47608868876161 | 17.344836089305563 | 11.770862699297004 | 12.440589232010563 | 2.0886481123932734 | 1.539759337462446 | 5.528833122537514 | 2.632693674246787 | 0.20872028241288265 | 1.1457445792816463 | 2.3816911775158807 | 0.4113981286859172 | 2.0301348760889146 |
| G | 45.289159275328664 | 19.58416309098553 | 11.09520561378354 | 8.724522163207013 | 2.39553023916804 | 2.2068723568315938 | 2.998859785176502 | 2.08833662434193 | 0.7138330063354663 | 0.7746962039154501 | 1.0933206243019327 | 0.8361270423925132 | 2.1993739742318255 |
| H | 44.00849531717684 | 18.674068712526598 | 11.673718202225489 | 9.50175628811375 | 2.808215794210352 | 2.252154383202021 | 2.6843734789246967 | 2.486272030639107 | 0.7463596846471823 | 0.7720382552307256 | 1.2106045693750276 | 0.7067618438100589 | 2.4751814399181513 |
| I | 45.95840963593769 | 16.63902446582157 | 13.804696841472214 | 9.964163616350659 | 2.392151225889026 | 1.8894003564473152 | 1.355566613671951 | 3.652988766170095 | 0.3652960332261899 | 0.9011592171996968 | 0.8570930320627942 | 0.3270463961008501 | 1.89300379964995 |
| J | 50.92386634777315 | 16.198921574751303 | 11.997285914901505 | 7.094145702923903 | 2.3153070477449735 | 1.7836808928240389 | 4.4124452250406385 | 1.0839841415266767 | 0.5768454301939894 | 0.6285612591290656 | 0.9200060359278311 | 0.4392756016228177 | 1.6256748256401046 |
###### Chart
| Category | p_Ascomycota | p_Basidiomycota | p_Mortierellomycota | p_Rozellomycota | others |
|---|---|---|---|---|---|
| A | 90.0 | 6.0 | 1.0 | 1.0 | 2.0 |
| B | 86.0 | 10.0 | 2.0 | 0.0 | 2.0 |
| C | 88.0 | 4.0 | 4.0 | 0.0 | 4.0 |
| D | 73.0 | 7.0 | 2.0 | 1.0 | 17.0 |
| E | 90.0 | 4.0 | 3.0 | 1.0 | 2.0 |
| F | 81.0 | 3.0 | 3.0 | 7.0 | 6.0 |
| G | 89.0 | 6.0 | 2.0 | 0.0 | 3.0 |
| H | 90.0 | 2.0 | 2.0 | 1.0 | 5.0 |
| I | 93.0 | 4.0 | 2.0 | 0.0 | 1.0 |
| J | 91.0 | 4.0 | 2.0 | 0.0 | 3.0 |
F
2.0
I
D
PC2 (18.1%)
J
C
0.0
G
E
H
B
-2.0
A
-2.5
0.0
2.5
5.0
7.5
PC1 (66.1%)
Suppl.Fig.1: Differences in chemical profile and taxonomic composition over the ten vineyards. (a) Plot of the two major Principal Components of chemical profile explaining in total 82.2% of the variation. (b) Taxonomic structure of Fungi in the vineyard soil based on ITS sequences. (c) Taxonomic structure of Bacteria and Archaea in the vineyard soil based on 16S sequences.

### Slide 2
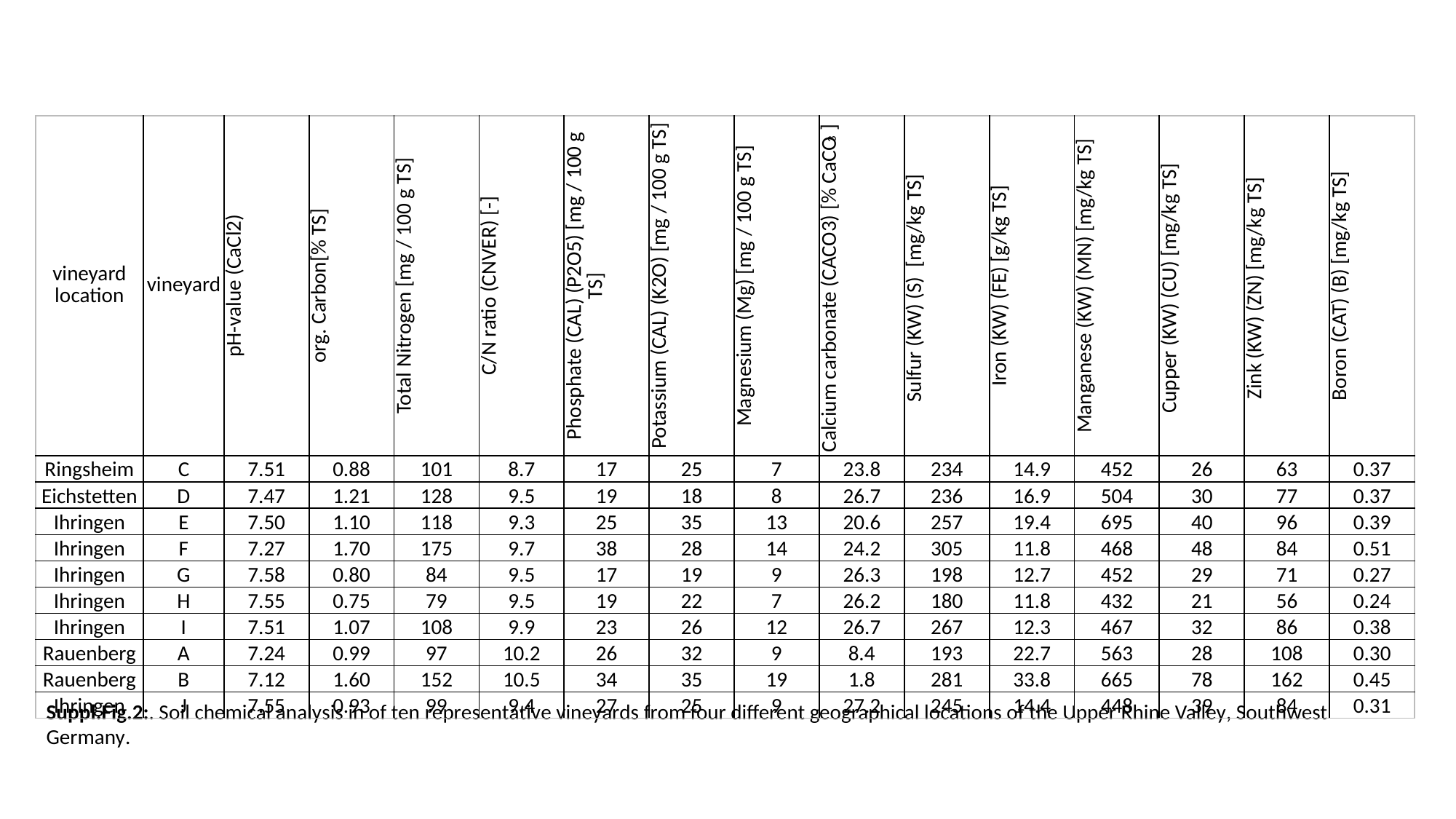

| vineyard location | vineyard | pH-value (CaCl2) | org. Carbon[% TS] | Total Nitrogen [mg / 100 g TS] | C/N ratio (CNVER) [-] | Phosphate (CAL) (P2O5) [mg / 100 g TS] | Potassium (CAL) (K2O) [mg / 100 g TS] | Magnesium (Mg) [mg / 100 g TS] | Calcium carbonate (CACO3) [% CaCO_3_] | Sulfur (KW) (S) [mg/kg TS] | Iron (KW) (FE) [g/kg TS] | Manganese (KW) (MN) [mg/kg TS] | Cupper (KW) (CU) [mg/kg TS] | Zink (KW) (ZN) [mg/kg TS] | Boron (CAT) (B) [mg/kg TS] |
| --- | --- | --- | --- | --- | --- | --- | --- | --- | --- | --- | --- | --- | --- | --- | --- |
| Ringsheim | C | 7.51 | 0.88 | 101 | 8.7 | 17 | 25 | 7 | 23.8 | 234 | 14.9 | 452 | 26 | 63 | 0.37 |
| Eichstetten | D | 7.47 | 1.21 | 128 | 9.5 | 19 | 18 | 8 | 26.7 | 236 | 16.9 | 504 | 30 | 77 | 0.37 |
| Ihringen | E | 7.50 | 1.10 | 118 | 9.3 | 25 | 35 | 13 | 20.6 | 257 | 19.4 | 695 | 40 | 96 | 0.39 |
| Ihringen | F | 7.27 | 1.70 | 175 | 9.7 | 38 | 28 | 14 | 24.2 | 305 | 11.8 | 468 | 48 | 84 | 0.51 |
| Ihringen | G | 7.58 | 0.80 | 84 | 9.5 | 17 | 19 | 9 | 26.3 | 198 | 12.7 | 452 | 29 | 71 | 0.27 |
| Ihringen | H | 7.55 | 0.75 | 79 | 9.5 | 19 | 22 | 7 | 26.2 | 180 | 11.8 | 432 | 21 | 56 | 0.24 |
| Ihringen | I | 7.51 | 1.07 | 108 | 9.9 | 23 | 26 | 12 | 26.7 | 267 | 12.3 | 467 | 32 | 86 | 0.38 |
| Rauenberg | A | 7.24 | 0.99 | 97 | 10.2 | 26 | 32 | 9 | 8.4 | 193 | 22.7 | 563 | 28 | 108 | 0.30 |
| Rauenberg | B | 7.12 | 1.60 | 152 | 10.5 | 34 | 35 | 19 | 1.8 | 281 | 33.8 | 665 | 78 | 162 | 0.45 |
| Ihringen | J | 7.55 | 0.93 | 99 | 9.4 | 27 | 25 | 9 | 27.2 | 245 | 14.4 | 448 | 39 | 84 | 0.31 |
Suppl.Fig.2:. Soil chemical analysis in of ten representative vineyards from four different geographical locations of the Upper Rhine Valley, Southwest Germany.

### Slide 3
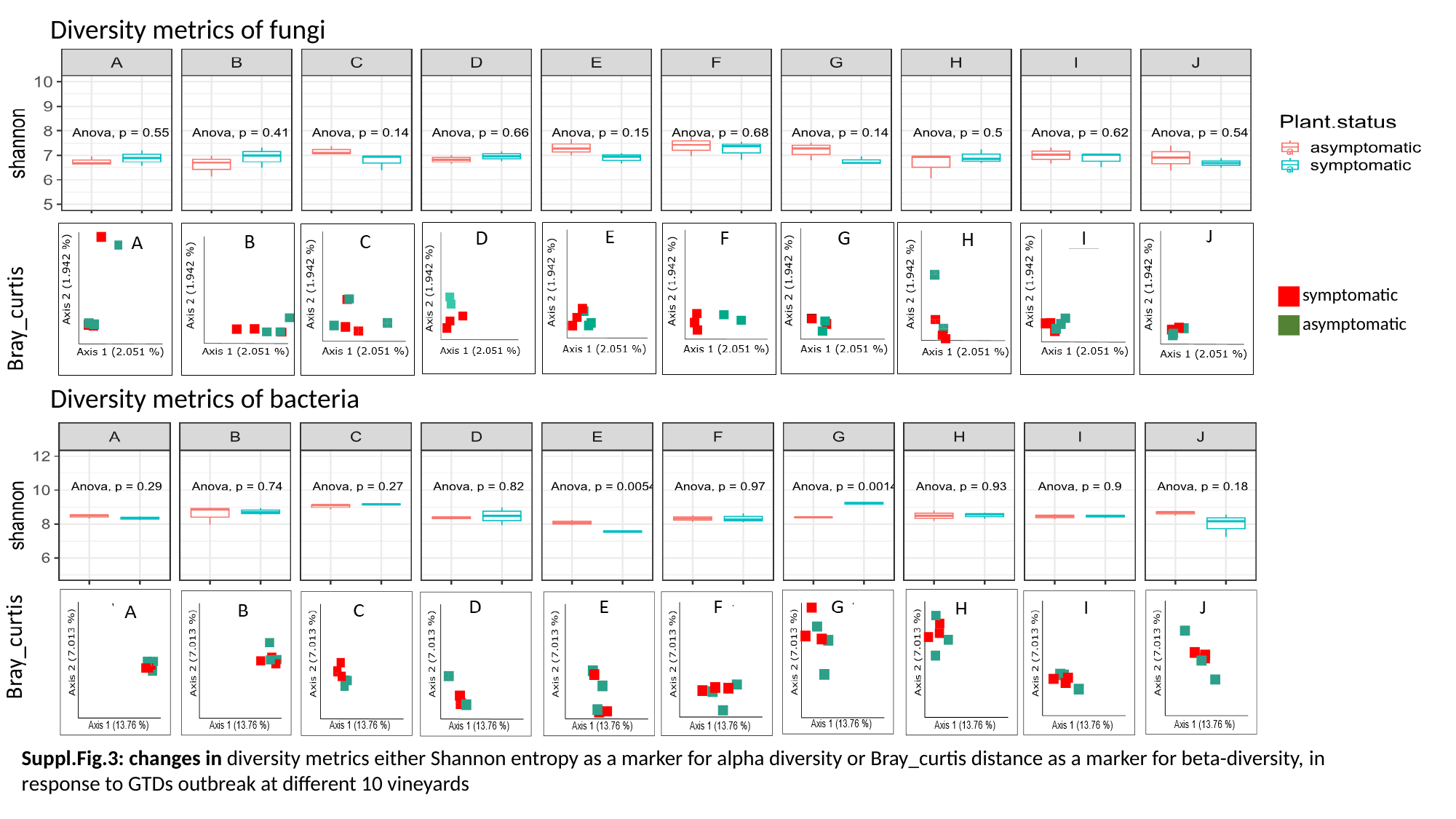

Diversity metrics of fungi
symptomatic
asymptomatic
Diversity metrics of bacteria
Suppl.Fig.3: changes in diversity metrics either Shannon entropy as a marker for alpha diversity or Bray_curtis distance as a marker for beta-diversity, in response to GTDs outbreak at different 10 vineyards

### Slide 4
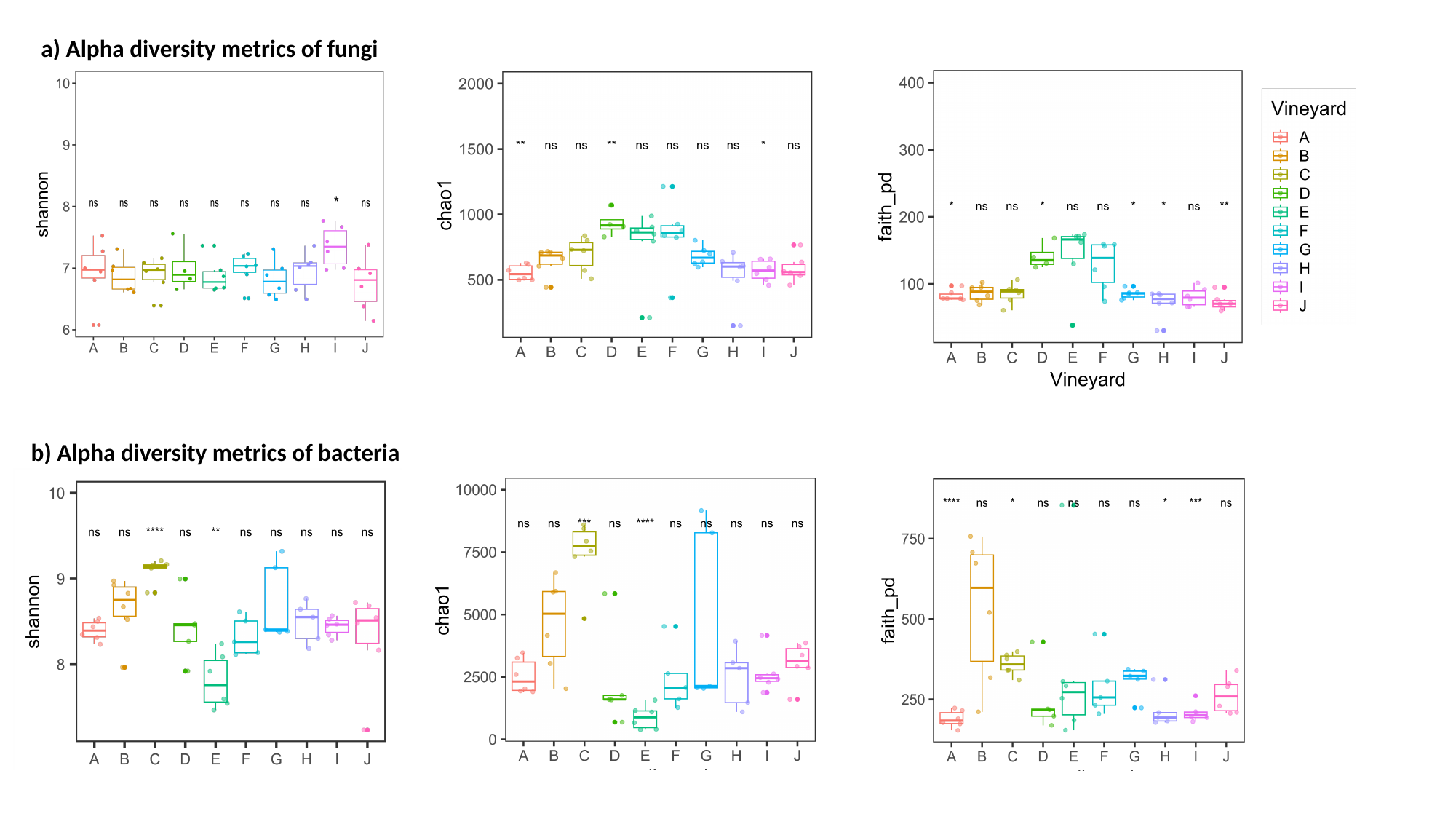

a) Alpha diversity metrics of fungi
b) Alpha diversity metrics of bacteria

### Slide 5
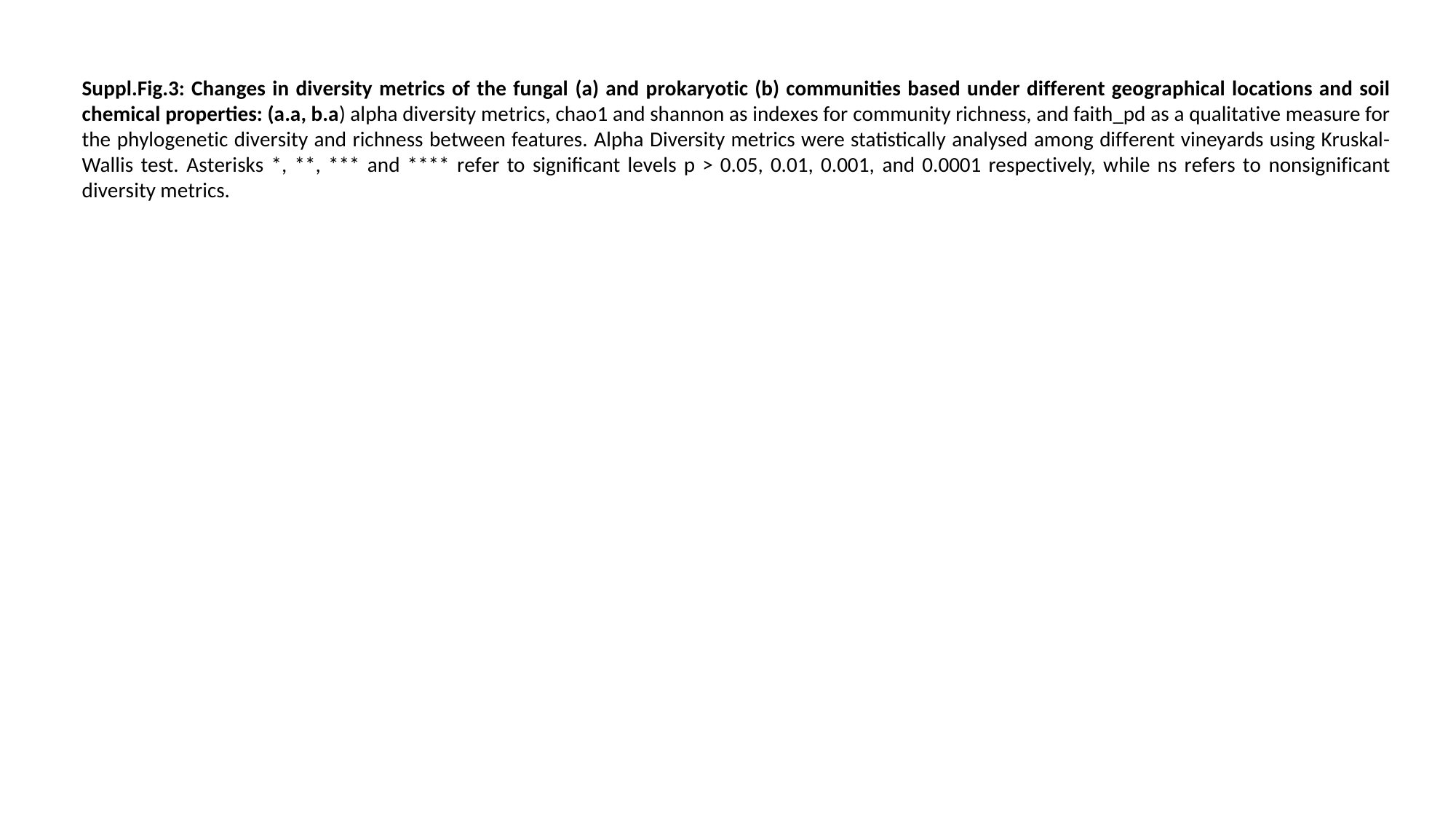

Suppl.Fig.3: Changes in diversity metrics of the fungal (a) and prokaryotic (b) communities based under different geographical locations and soil chemical properties: (a.a, b.a) alpha diversity metrics, chao1 and shannon as indexes for community richness, and faith_pd as a qualitative measure for the phylogenetic diversity and richness between features. Alpha Diversity metrics were statistically analysed among different vineyards using Kruskal-Wallis test. Asterisks *, **, *** and **** refer to significant levels p > 0.05, 0.01, 0.001, and 0.0001 respectively, while ns refers to nonsignificant diversity metrics.

### Slide 6
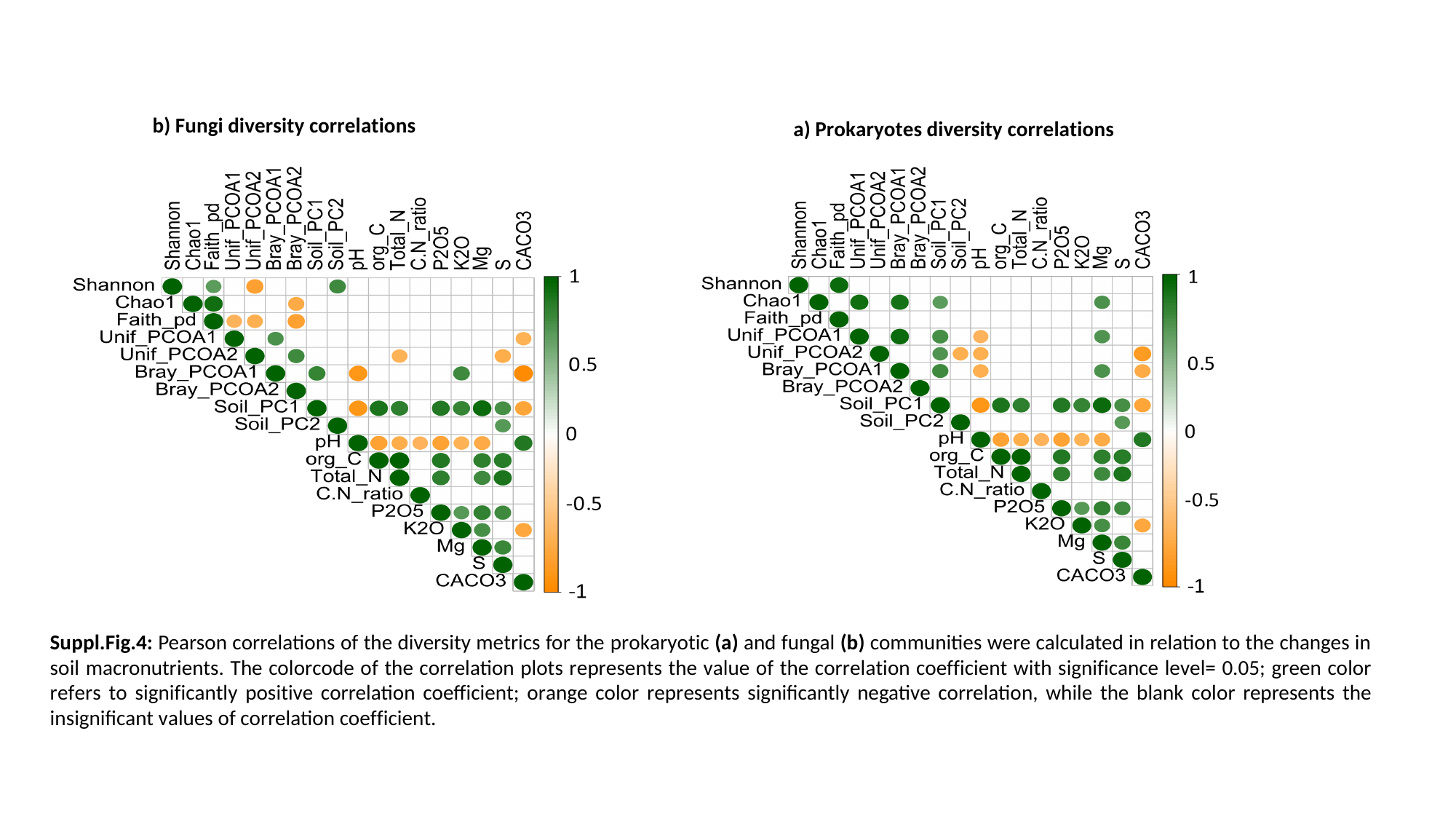

b) Fungi diversity correlations
a) Prokaryotes diversity correlations
Suppl.Fig.4: Pearson correlations of the diversity metrics for the prokaryotic (a) and fungal (b) communities were calculated in relation to the changes in soil macronutrients. The colorcode of the correlation plots represents the value of the correlation coefficient with significance level= 0.05; green color refers to significantly positive correlation coefficient; orange color represents significantly negative correlation, while the blank color represents the insignificant values of correlation coefficient.
